# Pathological stiffening by crosslinking glycation of titin

**DOI:** 10.64898/2026.02.13.705744

**Authors:** Agata Bak, Andra C. Dumitru, Alejandro Clemente-Manteca, Enrique Calvo, Roberto Silva-Rojas, Natalia Vicente, David Sánchez-Ortiz, Inés Martínez-Martín, Diana Velázquez-Carreras, Maria Rosaria Pricolo, Francisco M. Espinosa, João Almeida-Coelho, Carlos Pérez-Medina, Ricardo Garcia, Jesús Vázquez, Inês Falcão-Pires, Elías Herrero-Galán, Miquel Adrover, Jorge Alegre-Cebollada

**Author notes:** Equal contribution (Agata Bak, Andra C. Dumitru). Corresponding authors.,; X: @AlegreCebollada.

## Abstract

Heterogeneous, non-enzymatic glycation chemistry triggered by sugar-derived metabolites is typical of diseases that also entail pathological stiffening of cells, such as diabetes and age-related disorders. However, the mechanisms responsible for cell stiffening and the role of glycated biomolecules remain largely unexplored. Here, we show that glycation of cardiac titin, a giant intracellular protein scaffolding contractile sarcomeres, is increased in diabetes and leads to rigidification of both the protein and cardiomyocytes. Mechanistically, glycation-induced titin stiffening results from decreased contour length and enhanced folding of otherwise structurally intact protein domains following extensive formation of intramolecular crosslinking advanced glycation end products (AGEs). These stiffening effects outweigh softening contributions by competing, non-crosslinking AGEs. In combination, our work overcomes the intrinsic chemical complexity typical of glycation to uncover crosslinking AGEs as a source of pathological stiffening of cells, which we propose contributes to tissue dysfunction in situations of glycative stress.

## Introduction

Mechanics play inextricable roles in biological processes like development or stem cell differentiation^1^, and, when dysregulated, contribute to the onset and progression of disease. For instance, stiffening of the extracellular matrix (ECM) is linked to the growth of cancer cells^2^, and reduced compliance of the cardiac walls leads to heart failure because ventricles cannot properly accommodate blood during diastole^3^. Based on these observations, drugs countering mechanical alterations are gaining increasing traction in the clinic^4,5^. However, progress is hampered by an incomplete understanding of the molecular mechanisms underlying pathogenic changes in tissue mechanics. Here, we address why diabetic tissues and cells are stiffer than healthy counterparts, a situation that contributes to increased risk of heart failure among other systemic complications^6–8^ (**Figure 1A**).

**Figure 1.**
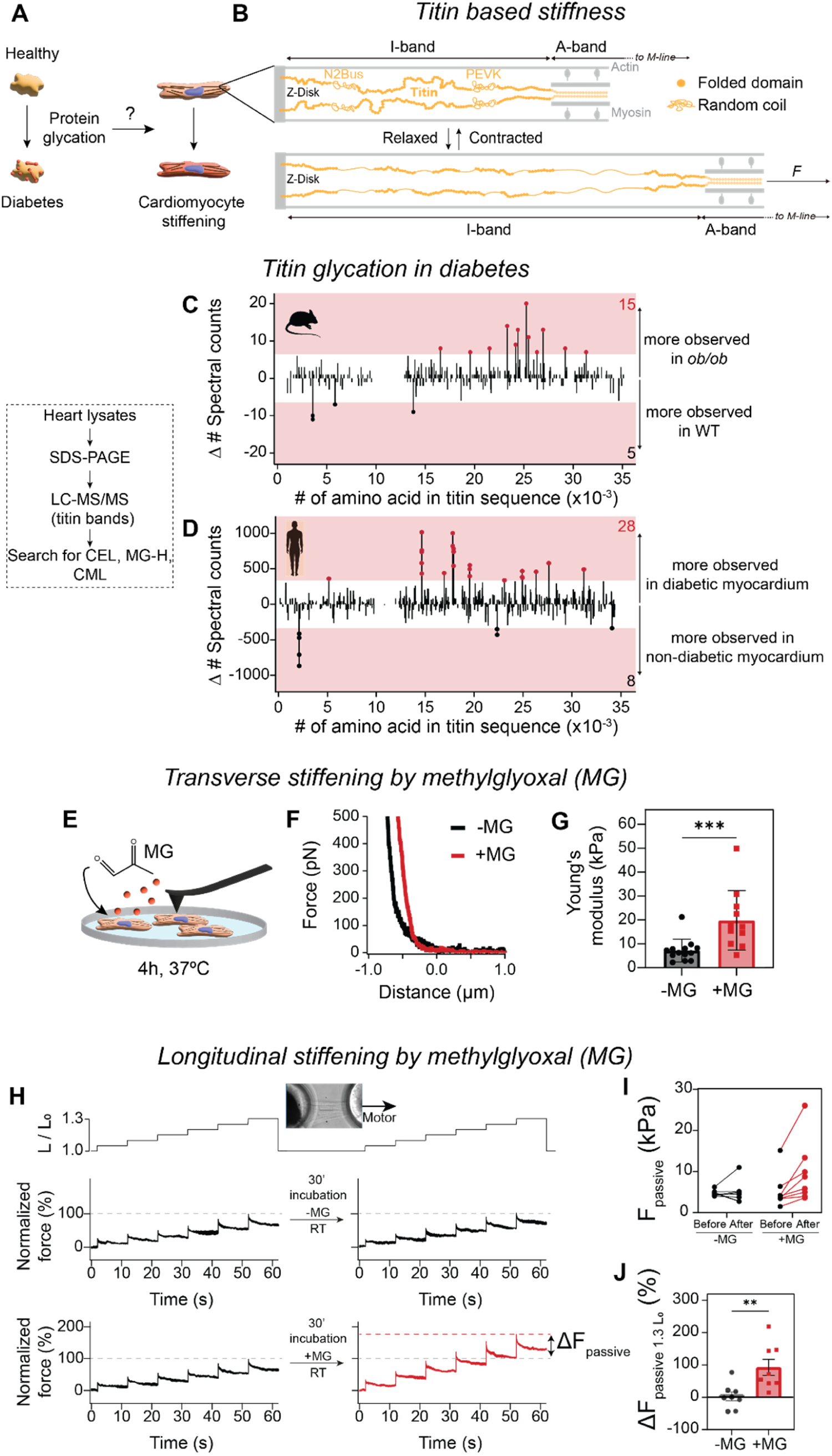
Glycation stiffens cardiomyocytes. **(A)** Hypothesis of this work: protein glycation contributes to cardiomyocyte stiffening in situation of glycative stress including diabetes. **(B)** Schematic representation of a contracted and relaxed I-band in half a sarcomere (not to scale). Titin is colored in yellow, while other sarcomeric proteins appear in gray. Titin Ig and fibronectin domains are represented as filled circles, and the approximate positions of the mostly unstructured N2Bus and PEVK segments are indicated. The length of the mechanically active I-band and the beginning of the A-band are delimited by arrows. Please note that in the relaxed sarcomere, unstructured regions are extended, and a fraction of Ig domains is unfolded. **(C)** Difference in spectral counts (Δ#Spectral counts) of peptides containing glycated residues along the titin sequence (x-axis) resulting from the hearts of 5 control and 5 *ob/ob* mice (2 chromatographic runs per sample). **(D**) Difference in spectral counts of peptides containing glycated residues between diabetic myocardium samples (n=2) and control myocardium samples (n=2). In C and D, the pink area indicates values above and below 3 x SD of control datasets where no differences in glycated peptides are expected (**Figure S1**). **(E)** Representation of AFM nanoindentation experiments to characterize the effects of incubation with MG on transverse stiffness of skinned neonatal cardiomyocytes. **(F)** Representative approach force-distance AFM curves recorded for untreated (-MG, black) and MG-treated (+MG, red) cardiomyocytes. **(G)** Quantification of Young’s moduli obtained by AFM nanoindentation. Each dot represents the median Young’s modulus obtained for an individual cell, results obtained with n = 13 for -MG condition and n = 11 for +MG condition, error bars represent SD. **(H)** Stretch protocol used to measure the passive force of cardiomyocytes in the longitudinal direction. A single skinned cardiomyocyte is mounted between a force sensor and a motor (inset). The cell is stretched from 1.0 to 1.3 times its initial length (L_0_) in 6 steps, and the passive force is recorded before and after 30-minute incubation with phosphocreatine-free relaxing buffer including or not MG at RT. Force traces show average results from n = 8 cells per condition; individual force traces were normalized considering the peak force values at 1.3 L_0_ before the incubation phase. **(I)** Changes in passive force measured at 1.3 initial length (L_0_) before and after incubation with and without MG. **(J)** Relative changes in passive stiffness after incubation with and without MG. Error bars represent SEM (n = 8). p-value <0.01(**), <0.001 (***).

The macroscopic mechanical properties of tissues derive from the nanomechanics of their constituent biomolecules. This is best exemplified in striated muscle, where both extra- and intracellular proteins are well-known to set tissue mechanics^9^. Key players include collagens^10^, the main structural scaffold of the ECM, and the giant intracellular protein titin^11–13^, a major contributor to cardiomyocyte (and myocardial) stiffness at physiological strains^9^ (**Figure 1B**). Considering this emergence across scales, it comes as no surprise that modulation of protein nanomechanics, for instance through protein isoform switch or by posttranslational modifications (PTMs), represents a physiological regulator of global tissue mechanics^11,14^. Importantly, in some diseases these very same mechanisms can result in pathological maladaptation perturbing tissue homeostasis. Such detrimental effects can be especially relevant in the case of PTMs favored during events of chemical stress, as exemplified by oxidations triggered by ischemia/reperfusion damage^15,16^. The mechanical consequences of other forms of chemical stress, including those induced by glycation typical of diabetes and age-related diseases^17,18^, remain incompletely understood. This so-called glycative stress results from abnormally high concentrations of sugars and/or related metabolites, which induce a complex cascade of non-enzymatic reactivity between their carbonyl groups and amino or guanidino groups of both extra- and intracellular proteins. These reactions eventually form a heterogeneous set of compounds known as advanced glycation end products (AGEs)^19^, which includes both crosslinking and non-crosslinking modifications. Most often, many different AGEs coexist in affected tissues, challenging integrative functional characterization^19^.

The main intracellular glycating compound is methylglyoxal (MG), a small and highly electrophilic α-oxoaldehyde formed as a side product of glycolysis^20,21^. MG is thousands of times more reactive than glucose^22^; thus, it can rapidly and extensively modify proteins even in the low μM concentration found in cells^22,23^. MG reacts with arginine side chains to form, among others, MG-derived hydroimidazolones (MG-H1, MG-H2 and MG-H3)^23^, and with lysine side chains, mainly yielding N^ε^-carboxyethyllysine (CEL)^24^ and a crosslinking MG-derived lysine dimer (MOLD)^25^.

Despite the concurrence of glycative stress and tissue stiffening, the causal relationship between them has remained elusive (**Figure 1A**). While crosslinking AGEs targeting ECM proteins have been proposed to contribute to tissue rigidification^6,26–28^, it remains unclear what the effect of competing, non-crosslinking counterparts may be since this type of modifications is expected to induce protein softening^29^. More importantly, mechanical alterations of the ECM cannot explain why diabetic or aged cells themselves are stiffer, as observed with endothelial cells^30,31^ and cardiomyocytes^6,32,33^. Building on circumstantial observations that titin is glycated in diabetic and aged myocardium^34,35^, we hypothesized that titin glycation could contribute to cardiomyocyte stiffening in these conditions. To test this hypothesis, here we have exploited single-molecule force spectroscopy, mass spectrometry, computer simulations, high-resolution NMR and cell mechanics experiments to demonstrate that AGEs in titin stiffen cardiomyocytes in situations of glycative stress.

## Results

### Increased glycation products in titin from diabetic myocardium

To study whether titin glycation plays a role in pathological cardiomyocyte stiffening, we first set out to provide additional support to available evidence indicating that cardiac titin glycation increases in conditions inducing glycative stress^34,35^. To this aim, we applied low-percentage sodium dodecyl sulfate-polyacrylamide electrophoresis (SDS-PAGE) to protein extracts obtained from the myocardium of 20-week-old *ob/ob* mice, a model of type II diabetes induced by obesity^36^, and from WT controls. Next, we sliced bands corresponding to titin and digested samples with chymotrypsin. Ensuing mass spectrometry (MS) analysis identifies three times more peptides containing CEL, MG-H and N^6^-carboxymethyllysine (CML) AGEs in *ob/ob* animals than controls (**Figures 1C, S1A**). Similarly, we find 3.5-fold enrichment in glycated titin peptides in human myocardium from type II diabetic patients compared to non-diabetic counterparts (**Figure 1D, S1B**).

### Cardiomyocyte stiffening by methylglyoxal

Our MS data confirm that pathological glycative stress during diabetes results in titin glycation, a modification that could contribute to cardiomyocyte stiffening^6,33^. To examine this possibility, we determined the passive stiffness of skinned cardiomyocytes undergoing glycative stress induced by MG. In a first approach, we used nanoindentation by atomic force microscopy (AFM), a method that measures transverse stiffness of specimens from force-distance (FD) curves that relate cantilever force to vertical displacement during indentation (**Figure 1E,F**). We find that typical FD curves recorded on neonatal murine cardiomyocytes treated with 50 mM MG for 4 h at 37°C have a steeper slope at the contact region compared with controls, indicative of MG-induced cell stiffening (**Figure 1F**). Quantification confirms a two-fold increase in the Young’s modulus of MG-treated cells (19±4 kPa, mean±SEM unless stated otherwise) compared to controls (7±2 kPa) (**Figure 1G**).

Next, we followed a complementary approach to assess the longitudinal stiffness of MG-treated cardiomyocytes. To this end, we tethered single skinned rat cardiomyocytes to a motor and a force transducer and applied a tensile test consisting of six stepwise length increases of 5% the initial length (L_0_). Each straining step causes a sharp rise in passive force, followed by a slow relaxation phase as expected from the viscoelastic nature of cardiomyocytes^29^. After this initial test, we incubated cells at slack length with buffer including or not 50 mM MG for 30 min at room temperature (RT), and the tensile test was repeated. To average data, we normalized passive force signals for each cell according to values at 1.3 L_0_ in the first tensile test (**Figure 1H**). Results indicate that MG treatment induces a 90% increase in passive force at 1.3 L_0_ from 5.0±0.8 kPa to 9.5±2.6 kPa, while incubations in the absence of MG show no effect (**Figure 1H-J**). Importantly, additional control experiments demonstrate that these stiffening effects do not result from potential MG-induced modification of the free amine group in ATP, a molecule present in the experimental buffers to prevent the formation of actin-myosin crossbridges^37^. In these control experiments, we find that relaxing buffer containing ATP preincubated with MG does not noticeably affect the passive stiffness of cardiomyocytes, in agreement with preserved blockage of actin-myosin interactions (**Figure S2**).

In combination, our cell mechanics experiments demonstrate that MG-induced glycation leads to cardiomyocyte stiffening in both transverse and longitudinal directions, in agreement with a recent report using skeletal muscle fibers^38^. Considering that titin is the main contributor to passive stiffness of cardiomyocytes, particularly in the longitudinal direction^39^, our data support that the increased glycation of cardiac titin in diabetes is indeed a contributor to cardiomyocyte rigidification. To mechanistically scrutinize this possibility, we examined the effects of MG-induced glycation on titin mechanics.

### Extensive formation of intradomain crosslinking AGEs in titin I91 domain following incubation with MG

The contribution of titin to cardiomyocyte stiffness results from the mechanical extension and contraction dynamics of the I-band domains of the protein^40^ (**Figure 1B**). Among them, the N2Bus and PEVK regions behave mostly as random coil/entropic springs that easily adapt their length to the pulling force in an elastic manner, while folded immunoglobulin-like (Ig) domains are more rigid because they sequester residues that do not contribute to the protein’s contour length. However, Ig domains can unfold upon application of force, which softens titin as a result of the associated increase in contour length (**Figure 1B**). This mechanical unfolding is reversible at forces lower than 10 pN^41^. Modulation of mechanical unfolding/folding transition kinetics in titin domains can result in large-scale changes in the mechanics of the full-length protein, typically with opposing outcomes for crosslinking and non-crosslinking modifications, as observed with oxidative modifications^29,42,43^.

To determine the mechanical effects of MG-induced glycation on titin Ig domains, we initially focused on the I91 Ig domain of the protein (also known as I27, PDB code 1TIT). I91 contains eight solvent-exposed lysine residues that can be modified by MG, potentially resulting in both crosslinking and non-crosslinking AGEs (**Figure 2A**). We first incubated purified (I91)_8_, a tandem repeat polyprotein containing eight domains of I91, in the presence of 50 mM MG for 72 h at 37°C and evaluated results using size-exclusion chromatography (SEC) (**Figures 2B,C, S3A**). We used two control samples, i.e. untreated, pristine (I91)_8,_ and (I91)_8_ incubated in the absence of MG. In the three conditions, we observe a peak at 15 mL elution volume marking monomeric (I91)_8_ (**Figures 2C, S3B**). Protein aggregation was evident in the sample incubated without MG, resulting in lower signal intensity for monomeric (I91)_8_ in this preparation. In contrast, protein aggregation was not detected in MG-treated (I91)_8_; in this case, the main (I91)_8_ peak is broader with higher contribution of species at lower elution volumes as expected from some degree of MG-induced intermolecular crosslinking. Unreacted MG was effectively separated from (I91)_8_, as indicated by the prominent peak appearing at the exclusion volume (20-22 mL). To examine the extent of modification by MG, we evaluated (I91)_8_ samples using MALDI-TOF/TOF mass spectrometry. While the spectrum of pristine (I91)_8_ shows a well-defined, narrow peak centered at 81.5 kDa matching the theoretical mass of the protein (81.7 kDa), the sample treated with MG yields a much broader peak at 87.6 kDa indicating the presence of a mixture of (I91)_8_ molecules containing different extents of MG-derived AGEs (**Figure 2D**). Accordingly, the band corresponding to MG-treated (I91)_8_ in SDS-PAGE gel is quite diffuse (**Figure S3B**).

**Figure 2.**
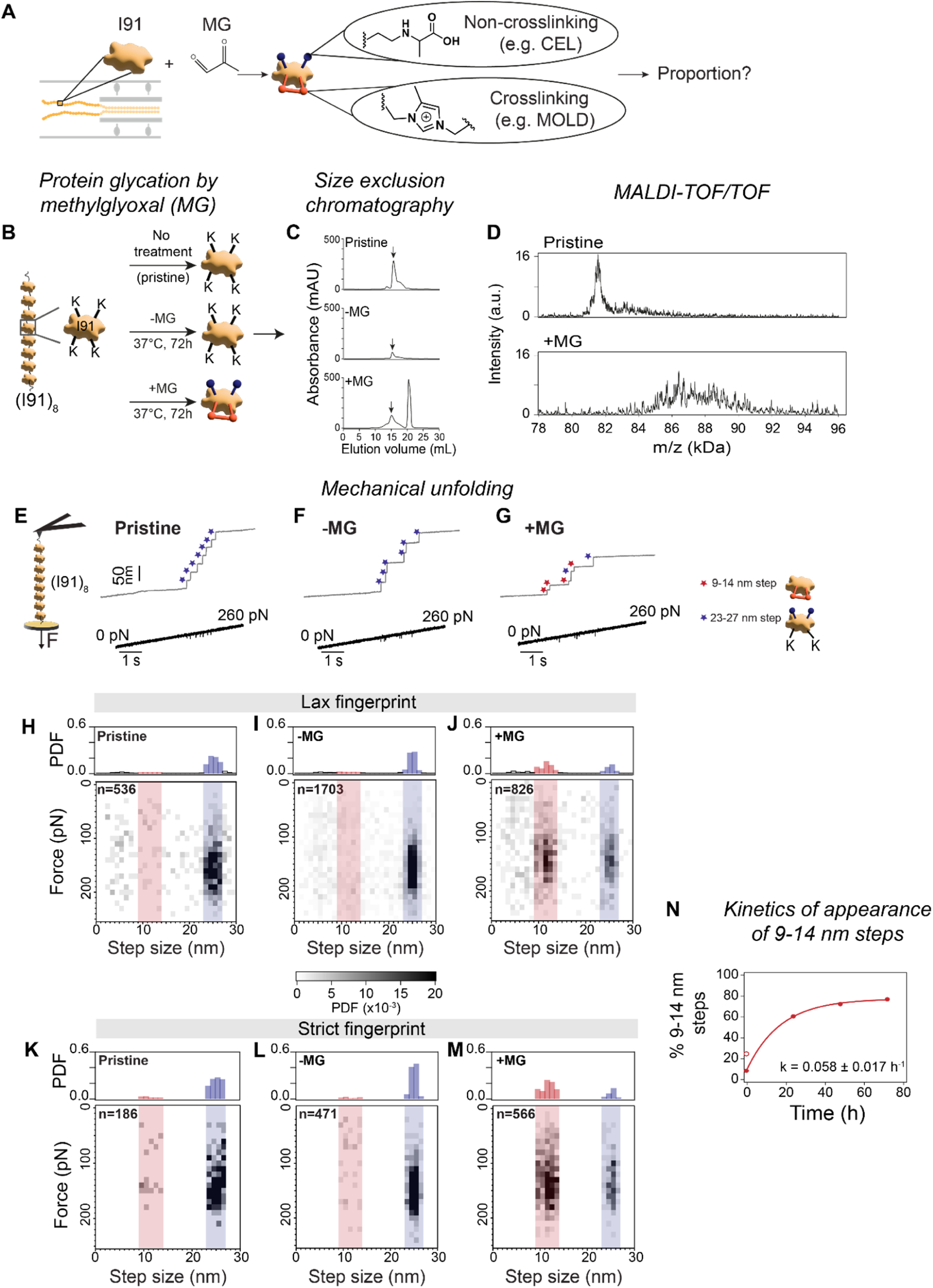
MG induces a high proportion of crosslinking modifications in the I91 domain of titin. **(A)** Representation of the reaction between I91 and MG leading to the formation of non-crosslinking (e.g. CEL; depicted in blue) and crosslinking (e.g. MOLD; depicted in red) AGEs in lysine residues. **(B)** Experimental groups to study the effects of glycation of the (I91)_8_ polyprotein. Non-modified lysine residues are represented in one-letter code; modifications indicated as in (A). **(C)** Size-exclusion chromatograms of (I91)_8_ protein preparations. Arrow marks elution volume of (I91)_8_. **(D)** MALDI-TOF/TOF spectra of pristine or glycated (I91)_8_ after incubation with 50 mM MG for 72 h at 37°C. **(E-G)** Representative single-molecule force-ramp traces to probe the mechanical unfolding of individual titin I91 domains in the three types of (I91)_8_ preparations. Blue stars mark 23-27 nm events coming from unfolding of full length I91 domains. Red stars indicate shorter events (9-14 nm). **(H-J)** Bidimensional histograms showing frequency of events according to their size and force at which they occur for the three experimental conditions, using lax fingerprinting to select single-molecule events. The number of events contributing to the data set are indicated. Monodimensional distributions of step sizes are shown on top of the bidimensional histograms. **(K-M)** Equivalent histograms to panels H-J obtained using strict fingerprinting to select single-molecule events. **(N)** Kinetics of appearance of 9-14 nm steps. The solid line is a fit to first order reaction kinetics. For the 0 h incubation time, the data for pristine protein was considered for the fit. The open symbol corresponds to the results from the sample to which MG was added and then immediately removed using FPLC. Error bars are SD of bootstrapping distributions^75^. PDF: probability density function.

Having proved that (I91)_8_ is extensively modified by MG, we used single-molecule force-spectroscopy by AFM (here referred to as AFS)^44^ to characterize the mechanical effects of glycation on I91. We subjected single (I91)_8_ proteins to a 40 pN s^-1^ increment in pulling force between 0-260 pN, while simultaneously measuring protein length (**Figure 2E-G**). As expected, mechanical unfolding of pristine (I91)_8_ or (I91)_8_ incubated in the absence of MG results in stepwise ∼25 nm increases in length at forces around 150-170 pN, where every step corresponds to a single I91 unfolding event (**Figure 2E,F**)^29^. Since our single-molecule experiments rely on non-specific physisorption of proteins to the AFM cantilever, we find a variable number of 25-nm events in individual recordings (seven and six in the examples in **Figure 2E,F**, respectively). Notably, many steps with shorter lengths are evident in single-molecule recordings obtained with the MG-treated (I91)_8_ sample (**Figure 2G**, short steps marked by red stars), suggesting that many unfolding domains contain covalent bonds that prevent full mechanical extension of the polypeptide, as previously observed for disulfide and isopeptide bonds^45,46^.

To quantify the extent of MG-derived crosslinking modifications in an unbiased manner, we selected single-molecule recordings following increasingly stringent fingerprinting criteria^44^. Initially, using a lax criterion, we extracted the step size and unfolding force of all events in traces containing ≥2 events with at least one being 25 nm in length, and represented the results in bidimensional histograms. For pristine (I91)_8_, this analysis identifies the expected main population of events centered at 25 nm that emerges from a background of non-specific events typical of single-protein AFS measurements^44^ (**Figure 2H**, 25 nm population shaded in blue). Similar results are obtained with control (I91)_8_ incubated in the absence of MG (**Figure 2I**). In contrast, the abundance of ∼25 nm events considerably drops in the sample treated with MG, where a new population of events between 9-14 nm is now observed (**Figure 2J**; population of short steps shaded in light red). For precise estimation of the proportion of these shorter steps, we followed a stricter fingerprint criterion analyzing only traces that contain 9-14 nm and/or 23-27 nm steps (**Figure 2K-M**). We find that 77% of events in the MG-treated (I91)_8_ sample correspond to short steps, strongly suggesting that most I91 domains in these conditions contain MG-induced covalent crosslinking modifications. By setting incubations with MG of different duration, we find that the proportion of short steps reaches saturation after 48 h of reaction (**Figures 2N, S4**). Importantly, these experiments also reveal that the sole addition of MG induces intradomain crosslinks in 24% of I91 domains despite its immediate removal by SEC, a process that is completed in less than 1 h at 4°C (open circle in **Figure 2N**). This result indicates that MG quickly induces a subset of covalent crosslinking modifications in the I91 domain of titin.

Overall, our single-molecule experiments with MG-treated (I91)_8_ indicate that intradomain covalent crosslinking modifications are very prevalent and can target the majority of I91 domains despite expected chemical competition with non-crosslinking modifications.

### Detection of crosslinking modification MOLD in MG-treated I91

We sought to characterize the chemical nature of the crosslinking AGEs formed in MG-treated I91. With that aim, we first produced glycated and control (I91)_1_ preparations (**Figures 3A, S5, Text S1**). Similar to (I91)_8_, the molecular mass of (I91)_1_ determined by MALDI-TOF/TOF increases upon 24 h incubation with 50 mM MG, and the peak corresponding to glycated (I91)_1_ is broader than that of the pristine protein (**Figure S6**). This indicates equivalent extent and heterogeneity of glycation in (I91)_1_ and (I91)_8_. Next, we collected the ^15^N,^1^H-TOCSY-HSQC NMR spectra of ^13^C,^15^N-(I91)_1_ samples. While the projection of the H-H planes in the spectrum for non-glycated (I91)_1_ displays well-defined cross-peaks at 7.2 ppm correlating lysine-H^ζ^ with the other intra-residue protons, these signals disappear in MG-treated (I91)_1_, which proves that MG modifies the NH^ζ^ group of the lysine side chains of (I91)_1_ (**Figure 3A,B**).

**Figure 3.**
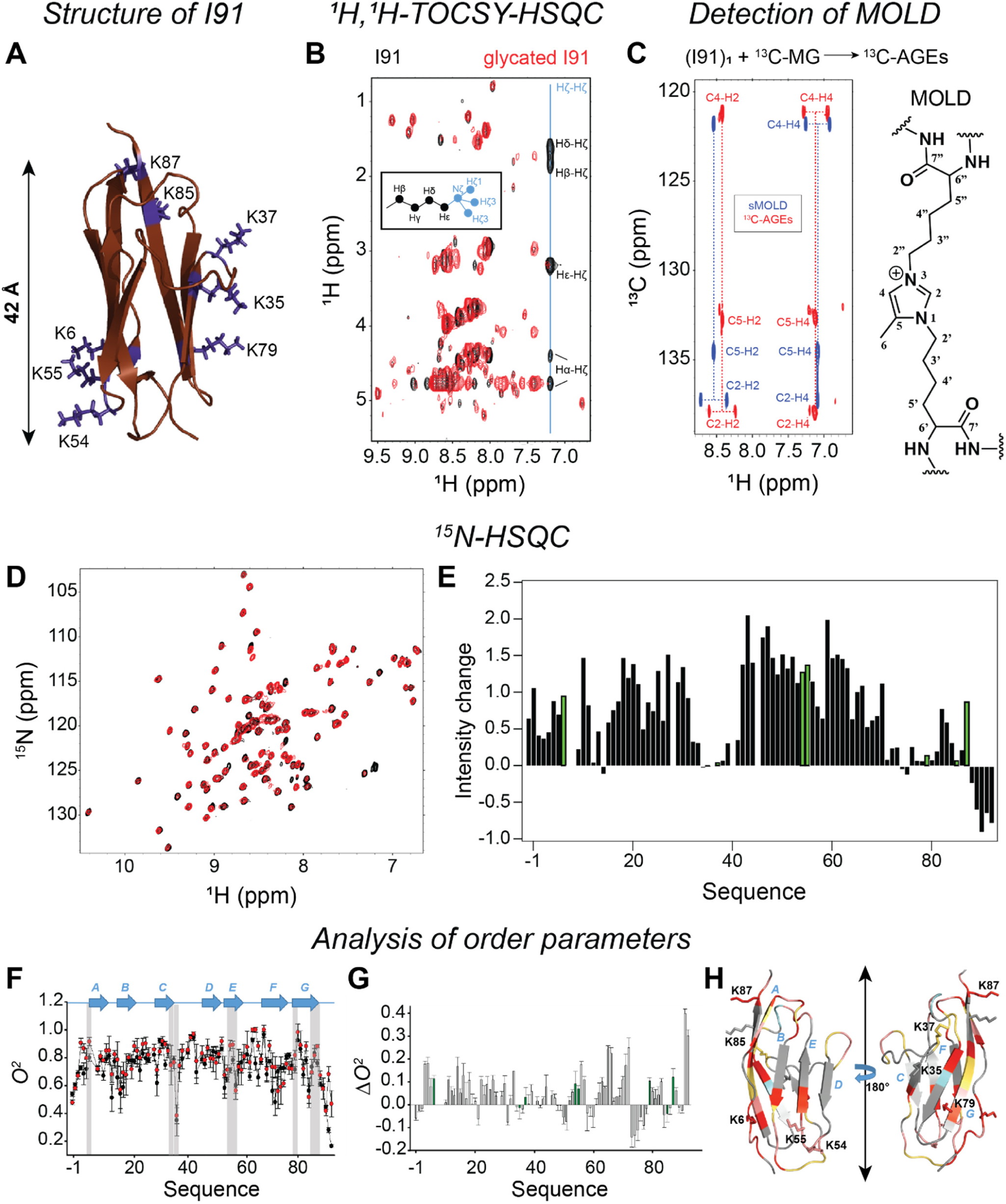
Characterization of glycation-induced modifications in titin’s I91 domain by NMR. **(A)** Ribbon representation of the solution structure of I91 (PDB: 1TIT). The side chains of lysine residues are shown in purple. The double arrow indicates the distance between the ends of the globular structure. **(B)** Overlapping of the projections of the different H-H planes of the ^1^H,^1^H-TOCSY-HSQC spectra obtained for native (black) and glycated (I91)_1_ (red). The chemical shifts of the H^ζ^-H^ζ^ cross-peaks corresponding to lysine residues are marked with a blue line that crosses the correlation peaks appearing due to coupling between the H_ζ_ with the other protons of the lysine side chains. The insert in the spectra shows a model representing the side chain of lysine. **(C)** Overlapping of the ^1^H,^13^C-HMBC spectrum obtained for sMOLD (blue) and that obtained for (I91)_1_ treated with ^13^C-MG (red). As it happens in ^1^H,^13^C-HMBC spectra, all the peaks corresponding to the C-H one bond correlation are split into two different signals separated by their ^1^J_C-H_ coupling constants. The chemical structure of MOLD is shown. **(D)** Overlay of the ¹⁵N-HSQC spectra for non-glycated (I91)_1_ (black) and glycated (I91)_1_ (red). **(E)** Effect of glycation on the intensity of the HSQC peaks, calculated as (I_glycated_/I_native_)-1, where I_native_ is the resonance intensity of each peak of I91 (I’_native_) internally corrected by the intensity of the N^δ^-H^δ^ cross peak corresponding to the side chain of N77 (see panel D) (I_native_=I’_native_/I_N77(Nδ-Hδ)_), and I_glycated_ is also the corrected resonance intensity of each peak in glycated (I91)_1_. The bars corresponding to lysine residues are colored in green. **(F)** Backbone *O*^2^ values obtained from NMR relaxation data for non-glycated (I91)_1_ (black) and glycated (I91)_1_ (red). Gray shaded areas indicate the positions of lysine residues. Position of β-strands following previously assigned annotations^81^ is shown as blue arrows at the top of the panel. **(G)** Differences between order parameters for residues in non-glycated (I91)_1_ and glycated (I91)_1_ (Δ*O*^2^= *O*^2^*_I91 glycated_* - *O*^2^*_I91_*). The bars corresponding to lysine residues are colored in green. **(H)** Cartoon representations of the 3D structure of I91, which have been color-coded according to the Δ*O*^2^ values plotted in panel **G** (blue for Δ*O*^2^ > 0.3, cyan for 0.2 < Δ*O*^2^ ≤ 0.3, red for 0.1 < Δ*O*^2^ ≤ 0.2, salmon for 0.05 < Δ*O*^2^ ≤ 0.1, and yellow for 0.02 < Δ*O*^2^ ≤ 0.05. The image presents two views of the same structure, rotated 180°. Labels for the β-sheets are shown in blue. Images were generated with Pymol^88^.

To investigate the chemical nature of the AGEs formed in glycated I91, we incubated unlabeled (I91)_1_ with ^13^C-labeled MG. The advantage of this strategy is that only carbons coming from MG become NMR-visible, thus allowing straightforward comparison with commercially available chemical standards of AGEs. The ^1^H,^13^C-HMBC spectra of unlabeled (I91)_1_ modified with ^13^C-MG evidence the correlation of some aromatic (i.e. between 6 and 9 ppm in the ^1^H dimension) one bond C-H cross-peaks with other C-H cross-peaks. This demonstrates that MG induces the formation of aromatic AGEs on I91 (**Figure 3C**). To assess whether these signals could arise from MOLD, we collected the ^1^H,^13^C-HMBC spectrum of MOLD standard (sMOLD) and carried out its chemical shift assignment (**Table S1**). The ^1^H,^13^C-HMBC signals obtained for sMOLD match those observed for glycated (I91)_1_ (**Figure 3C**), unequivocally demonstrating the formation of MOLD on MG-treated I91. Subtle shifts in the ^1^H,^13^C-cross-peaks of MOLD formed on I91 compared to sMOLD are likely due to the different chemical environments. In addition, the low intensity peaks appearing next to the C5-H2, C5-H4 and C2-H4 cross-peaks suggest that ^13^C-MG-treated (I91)_1_ might contain at least two different MOLD moieties with different protein chemical environments.

In combination, our NMR data confirm that modification of I91 lysines by MG is substantial and identify MOLD as a crosslinking AGE present in the modified protein.

### Restricted conformational flexibility and preserved fold of glycated I91

Our interpretation of the AFS data assumes that glycated I91 remains folded in the absence of force application. To obtain independent validation of this assumption, we further used NMR spectroscopy to get insights at the residue level on potential structural effects in glycated I91. We observe that the positions of the structural fingerprint ^15^N-HSQC amide cross-peaks in (I91)_1_ do not change upon incubation with MG (**Figure 3D**), a first indication that glycation does not induce major alterations to the structure of I91. Furthermore, we do not detect any relevant change in the ring shifted resonance ^1^H signal of L58-^δ^CH_3_, which arises from its structural proximity to the aromatic side chain of W34 in the hydrophobic core of the domain (**Figure S7A,B**), nor in the long-range NOE signals that result from tertiary structural contacts (**Figure S7C**). Using the backbone chemical shifts (i.e. N, H_N_, C_α_, C_β_, H_α_ and CO) assigned for all residues in ^13^C,^15^N-(I91)_1_ incubated in the presence or absence of MG (BMRB codes 53390 and 53389, respectively), we also observe that glycation does not affect the β-sheet propensity scores of the β-sheets of the domain (**Figure S7D-F**). Altogether, these data prove that glycation mediated by MG does not impact the secondary nor the tertiary structure of I91.

Despite preserving the global fold of the I91 domain, MG-treated samples show increased intensity of many HSQC peaks (**Figure 3E**). This suggests that glycation shifts I91 towards a predominant conformation, thereby reducing exchange between minor conformational populations. The most affected residues are those mainly located at the ABED β-sheet, as well as in the loops connecting these strands and in those connecting them with the A’GFC β-sheet (**Figure S8A**). Intriguingly, the HSQC peaks that mostly change their intensities upon glycation are located at the N-terminus of strand D and at the C-terminus of strand E, which could indicate modification of nearby C47 and C63 (**Figure S8A**). However, the chemical shifts of the C_β_ of C47 and C63 (highly sensitive to the oxidation state of the thiol group) did not remarkably change upon glycation (27.3 ppm for C47 and 28.1 ppm for C63, **Figure S8B**). This observation aligns with previous results, which show that both cysteine residues in I91 have low solvent accessibility when the domain is folded, and therefore cannot be targeted by reagents in the solution at 37°C^29^ (**Figure S8C)**.

To evaluate the effect of glycation on the dynamics of I91, we acquired NMR relaxation data (i.e. R1, R2 and ^15^N HET-NOE; **Figure S9**) and determined the amplitudes of the conformational fluctuations of the backbone amide groups in terms of order parameters (*O*^2^, on the ps to ns time scale) according to a model-free formalism^47^. *O*^2^ values were calculated considering that the best-fit rotational diffusion tensor is anisotropic with a correlation time (τ_c_) of 10.6 and 11.5 ns for untreated and MG-treated (I91)_1_, respectively (**Figure 3F**). The mean values (±SEM) of the backbone order parameters <*O*^2^> calculated for the L1-L89 stretch (**Text S1**) are 0.701±0.011 for I91incubated in the absence of MG and 0.768±0.013 for MG-treated I91 (p<0.0001 for paired samples t-test), suggesting that glycation restricts overall protein conformational dynamics and rigidifies the domain. This result is even clearer from the comparison of *O*^2^ (i.e. Δ*O*^2^= *O*^2^*_I91 glycated_* - *O*^2^) at the residue level (**Figure 3G,H**). Remarkably, not all regions showing increased *O*^2^ are located near MG-sensitive lysine residues, which indicates that glycation can reduce structural fluctuations in an allosteric manner. Finally, the restricted conformational flexibility of glycated I91 was also captured by a reduced conformational entropy of the protein backbone (i.e. ΔS_conf_=-114.2±2.9 Jꞏmol^-1^ꞏK^-1^ or TΔS_conf_=-34±0.8 kJꞏmol^-1^ at 298 K; error is SD), as calculated from NMR relaxation data^48^.

In short, structural characterization of glycated I91 proves that MG does not induce any major change to the fold of the protein. However, we detect reduced conformational flexibility of several regions of the glycated domain, which probably results from the formation of crosslinking AGEs captured in single-molecule force-spectroscopy and NMR experiments.

### Mechanically relevant glycated lysines in I91

Our results so far indicate that MG-induced glycation of I91 leads to the formation of intradomain lysine-lysine crosslinking AGEs in the absence of major structural alterations. To directly examine the extent of this type of AGEs, we did additional AFS experiments using MG-treated (I91)_8_, but now including a protein preparation where we aimed to transform the lysine residues into inert CEL^49^ (**Figures 4A,B, S10**). Western blot (WB) analysis confirms that our experimental protocol results in high levels of CEL, which are considerably higher than in I91 incubated only with MG (**Figure 4C**). AFS results show that control samples not subjected to CEL modification but incubated with MG display the expected increase in short unfolding steps after MG incubation (**Figure 4D,E**). However, the appearance of short unfolding steps upon MG incubation is notably reduced in the case of the CEL-modified (I91)_8_ preparation (**Figure 4D-F**). Hence, this set of experiments confirm that lysine-lysine covalent crosslinking AGEs are responsible for the reduced mechanical unfolding length of glycated I91 domains.

**Figure 4.**
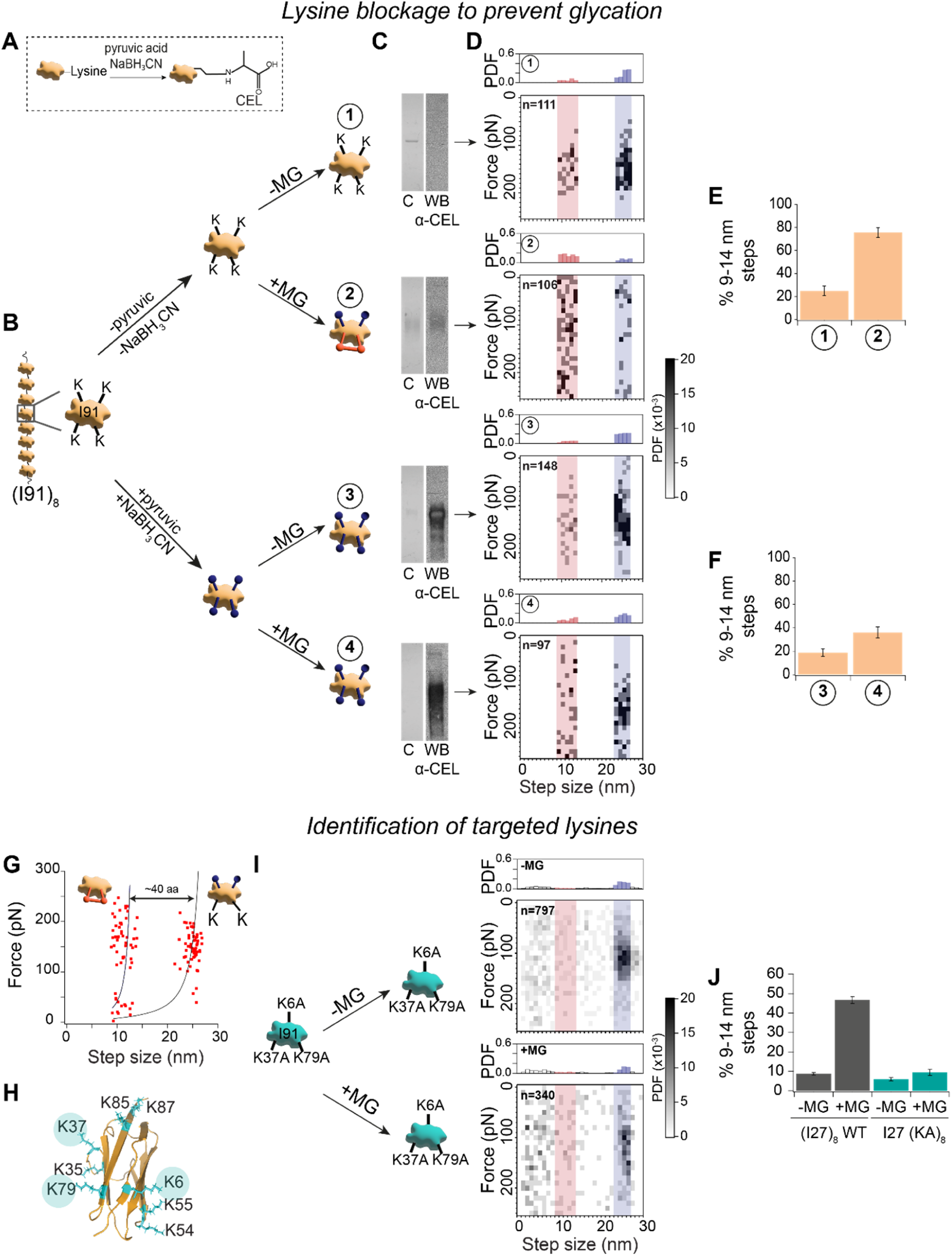
Lysines in I91 are mechanically relevant glycation targets. **(A)** Schematic representation of a lysine residue reacting with pyruvic acid in the presence of NaBH_3_CN to form CEL. **(B)** Experimental workflow to block lysines by formation of CEL (routes 3 and 4), which prevents subsequent MG glycation in route 4. Control incubations of the (I91)_8_ are also indicated (routes 1 and 2). **(C)** SDS-PAGE analysis of (I91)_8_ domains that have undergone the reactions shown in (B). Panel shows Coomassie blue staining (*left*) and WB analysis using anti-CEL antibodies (*right*). **(D)** Bidimensional histograms showing the frequency of the size of the unfolding steps and the force at which they occur in AFS traces selected with stringent fingerprinting for the four experimental conditions. The number of events contributing to the data set are indicated. Monodimensional distributions of step sizes are shown on top of the bidimensional histograms. **(E-F)** Quantification of the percentage of unfolding steps between 9 and 14 nm in the two experimental arms. **(G)** Distribution of force versus step size for events obtained using MG-incubated (I91)_8_ in AFS experiments. Solid lines are worm-like chain fits to the data. The contour length reduction in short steps is estimated to correspond to a covalent barrier trapping around 40 amino acids. **(H)** Structure of the I91 domain (PDB: 1TIT) highlighting lysine residues mutated out in (I91-KA)_8_. Image was generated with Pymol^88^. **(I-J)** (I91-KA)_8_ was incubated in the absence or presence of 50 mM MG for 72 h at 37°C and probed using single-molecule AFS. Single-molecule traces were selected with the non-stringent fingerprinting criterion, and data are represented as in panels D and E. WT data from Figure 2. Error bars are SD of bootstrapping distributions^75^. PDF: probability density function.

Next, we investigated the specific lysine pairs that are preferential crosslinking sites in MG-treated I91. We first analyzed the step sizes and associated forces resulting from mechanical unfolding of glycated I91. By fitting the worm-like chain model of polymer elasticity^50^ to the force *versus* step size distribution, we determined that the reduction in contour length of the I91 domain containing crosslinking AGEs is ∼16 nm (i.e. around 40 amino acids trapped by the covalent crosslink considering a contour length of 0.4 nm per amino acid^51^) (**Figure 4G**). We identified three candidate pairs of structurally vicinal lysine residues in the sequence of I91 that could account for this reduction in contour length, i.e. K6-K55, K35-K79 and K37-K85 (**Figure 4H**). To prove that MG-induced crosslinking involves at least one of these lysine pairs, we produced the mutant polyprotein (I91-KA)_8_, in which we replaced K6, K37 and K79 with non-reactive alanine residues in all I91 domains. Upon 72 h incubation of (I91-KA)_8_ at 37°C with 50 mM MG and subsequent purification (**Figure S11**), we conducted unfolding experiments by AFS. Different from the wild-type protein, the population of 9-13 nm steps in MG-treated (I91-KA)_8_ is only marginally increased with respect to the mutant protein incubated in the absence of MG (**Figure 4I,J**). In combination, our results with (I91-KA)_8_ confirm that at least one of the selected lysine pairs is a primary target for MG-induced crosslinking in I91. In agreement with this result, following an MS-based search for crosslinked peptides between lysine pairs in (I91)_1_ incubated with 50 mM MG for 24 h at 37°C, we were able to unambiguously detect the presence of a MOLD-crosslinked dipeptide involving K35 and K79 (**Figure 5**).

**Figure 5:**
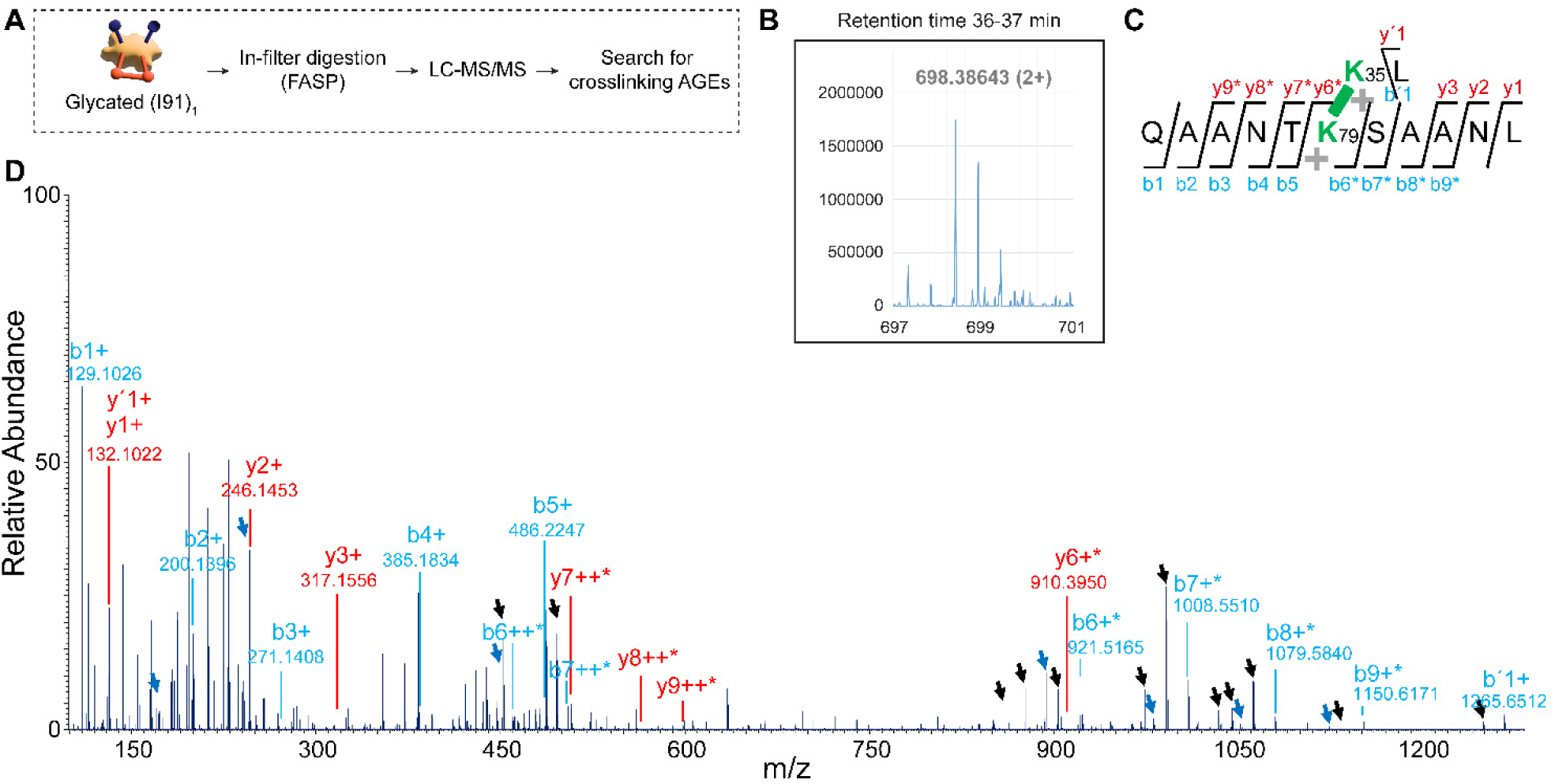
Identification of a MOLD crosslink in glycated I91 by LC-MS/MS. (**A**) Schematic representation of the experimental workflow. (**B**) Zoomed-in integrated survey scan between minutes 36 and 37, showing the doubly-charged precursor ion at m/z 698.38 corresponding to a chymotryptic dipeptide from I91 crosslinked by a MOLD connector. (**C**) Sequence of the chymotryptic dipeptide; the connector and lysines 35 and 79 are highlighted in green. (**D**) Assigned fragmentation MS/MS spectrum corresponding to the peptide in (C), taking the long peptide sequence (QAANTKSAANL) as a base on which the short sequence plus the crosslinking agent (MOLD-KL) are added. The figure shows the main fragmentation series (b and y) for the fragments singly- (+) or doubly-charged (++), containing (asterisks) or not the MOLD-KL delta-mass, demonstrating the exact location of crosslinking. The ý1 and b’1 ions would correspond to the breaking of the peptide bond between the residues of the short sequence (KL), giving rise to the common fragment derived from leucine (y1 / ý1) and the fragment corresponding to the rest of the crosslinked structure (b’1). Black arrows indicate neutral losses of water or ammonium. Blue arrows mark the a-series ions derived from the neutral loss of the carbonyl group from the corresponding b-series ions, demonstrating the correct assignment of these ions.

### Interdomain crosslinking AGEs targeting titin domains

In addition to intradomain crosslinking of lysine residues, interdomain and intermolecular crosslinking reactions should also be possible when MG reacts with serially linked protein domains such as those in (I91)_8_ (**Figure S12A**). While our preparations are devoid of high-molecular weight intermolecular crosslinks thanks to the size-exclusion purification step (**Figure S3B**), we cannot rule out contributions of intramolecular, interdomain crosslinks that would reduce the number of mechanically unfoldable domains in AFS experiments (**Figure S12B**). Indeed, we detect a slight decrease in unfolding events per AFS trace, from an average of ∼5 in controls to ∼4 in glycated (I91)_8_ (**Figure S12C**), suggesting that MG incubation induces some interdomain crosslinking. Considering that interdomain crosslinks would also cause structural modifications on the (I91)_8_ protein, we studied the impact of MG treatment on the overall structure and size of (I91)_8_ by small-angle X-ray scattering (SAXS). The scattering signals for both glycated and non-glycated (I91)_8_ overlap to a great extent, suggesting that the influence of glycation on the overall structure of (I91)_8_ is minor (**Figure S12D**). In addition, neither Porod-Debye plot shows a Porod plateau (**Figure S12E**), proving that neither protein preparation displays compact globular structures and that both have a certain degree of flexibility. This observation agrees with the plateau observed in the *q*^3^*ꞏI(q)* vs. *q*^3^ plots (**Figure S12F)** which is typical of flexible partially folded structures^52^. Moreover, both SAXS-derived pair-distance distribution functions (*P*r; probability distribution of the interatomic vectors inside the molecule) exhibit a local maximum at ∼20Å, which accounts for the inter-atomic distances within a single I91 domain, and another one at ∼46 Å, which indicates the separation between the mass centers of two consecutive I91 domains (**Figure S12G)**. This separation agrees with the average length of the I91 domain (**Figure 3A**), thus indicating extended domain organizations.

In summary, AFS and SAXS data taken together point to limited extent of interdomain crosslinking in MG-treated (I91)_8_ in our experimental conditions.

### Crosslinking glycation products in full-length titin

Since MG-induced glycation results in the formation of crosslinking AGEs in I91, we hypothesized that other titin domains could also be target of these modifications. To test this possibility, we examined by AFS and HaloTag technology^53^ the consequences of MG-induced glycation of native cardiac titin isolated from mice. In these experiments, we covalently attached engineered titin molecules containing a HaloTag inserted in the distal I-band region of the protein to glass surfaces coated with HaloTag ligand (**Figure 6A**). Next, we added 50 mM MG to the fluid chamber of the AFS and started pulling from titin molecules in constant-velocity mode for several hours. We also ran experiments in the absence of MG. Two representative force-distance traces are shown in **Figure 6A**; both of them exhibit a characteristic sawtooth pattern, a well-known single-molecule fingerprint in constant-velocity force spectroscopy where every peak originates from the mechanical unfolding of a titin domain and the distance between peaks reflects the increase in contour length associated with unfolding^54^. Using the worm like chain model, we find that the commonest change in contour length between consecutive peaks in the absence of MG is ∼30 nm (**Figure 6B**), as expected for non-modified titin domains^53^. In agreement with previous reports, we also detect a less frequent population of unfolding transitions (less than 20% of total events) characterized by changes in contour length shorter than 20 nm (**Figure B**)^42,53^. Interestingly, the distribution of unfolding events shifts towards shorter changes in contour length when titin is incubated with MG (**Figure 6A,B**), indicating that many domains in titin in addition to I91 undergo MG-induced crosslinking.

**Figure 6.**
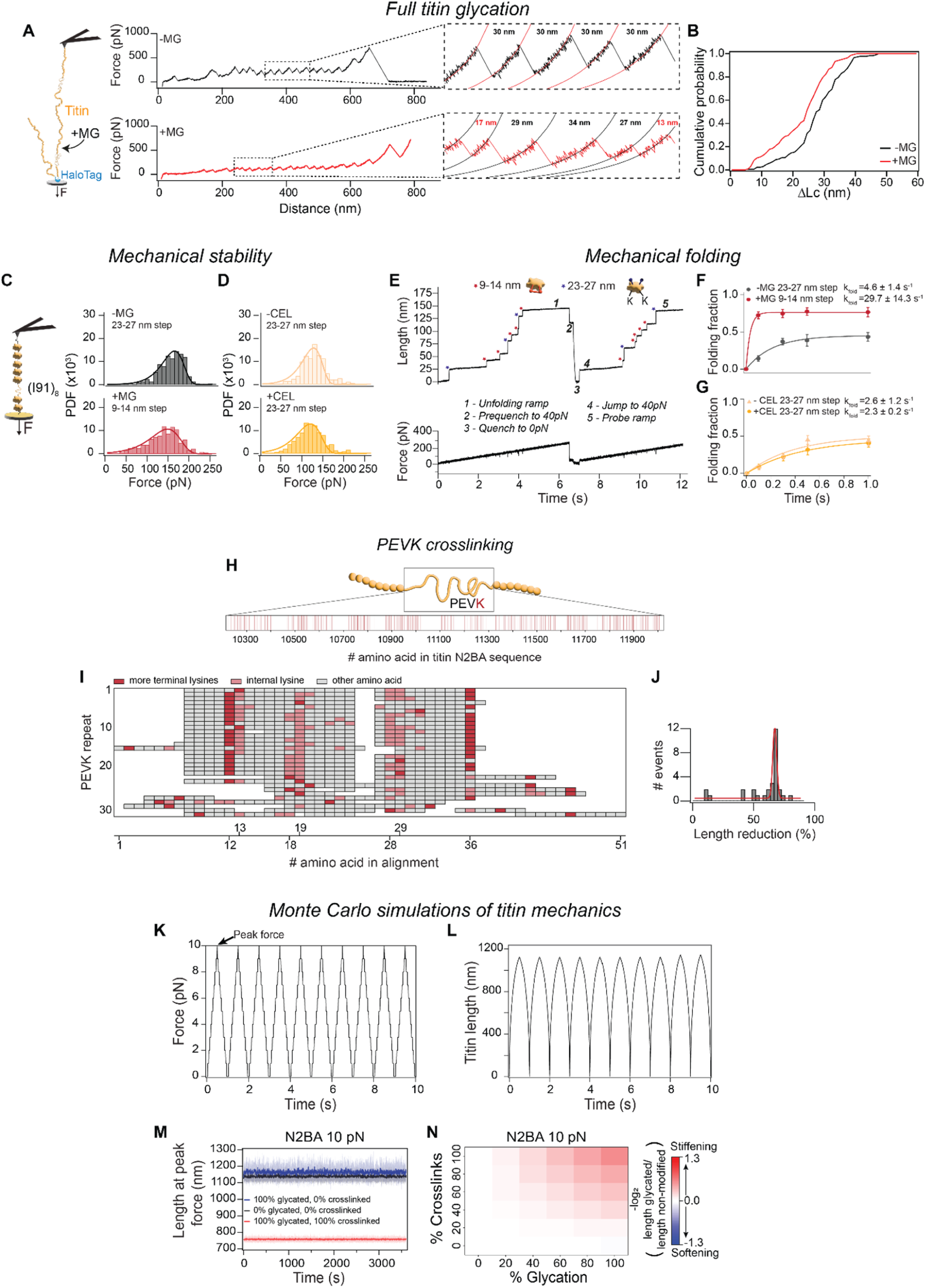
Mechanics of glycated titin. **(A)** Native titin molecules containing a HaloTag insertion at the end of the I-band region are immobilized on a HaloTag-ligand derivatized glass surface and stretched using AFM in constant-velocity mode. Glycation is induced *in situ* by adding 50 mM MG. Representative curves are shown. **(B)** Cumulative distribution of changes in contour length (ΔLc). **(C)** Distribution of unfolding forces of I91 domains after 72 h incubation of (I91)_8_ at 37°C in the absence (*top*; n = 440 events from 3 independent experiments) and in the presence of 50 mM MG (*bottom*; n = 436 9-14 nm events from 2 independent experiments). **(D)** Distribution of unfolding forces for (I91)_8_ incubated for 48 h at 50 °C without (*top*; n = 323 from 4 independent experiments) or with (*bottom*; n = 418 from 4 independent experiments) pyruvic acid and NaBH_3_CN. Solid lines in panels E and F are fits to the Bell-Evans model of force-activated reactions ^55,74^. **(E)** Representative unfolding-quench-probe AFS trace to study mechanical folding of glycated I91. A single glycated (I91)_8_ polyprotein is subjected to 40 pN s^-1^ unfolding pulse, then force is quenched to 0 pN and finally increased again in a probe pulse. The folding fraction is calculated by comparing the number of unfolded domains in both ramps. In this example, 5 out of 5 9-14 nm steps and 2 out of 3 23-27 nm steps refolded during the force quench. 40 pN force pulses are included for optimal sensitivity (see main text). **(F,G)** Folding fractions corresponding to the same protein preparations as in (C,D). Lines represent exponential fits to the data. In all cases, n>60 for each time point. **(H)** Schematic representation of the PEVK region in N2BA titin, highlighting the position of lysine residues in the sequence of human N2BA titin (Uniprot: Q8WZ42). **(I)** Alignment of the 31 PEVK repeats of human N2BA titin (repeats according to Uniprot: Q8WZ42). The most terminal lysine residues in each repeat are highlighted in red, while the remaining lysine residues are marked in pink. All other amino acids are shown in gray. **(J)** Distribution of contour length reduction in all PEVK repeats when their first and last lysine residues are crosslinked by MOLD. **(K)** Force protocol for Monte Carlo simulations. **(L)** Corresponding protein length of a virtual N2BA titin subjected to to the force protocol in (K). **(M)** Length of N2BA titin at 10 pN during the Monte Carlo simulations for non-modified (black), 100% glycated/0% crosslinked (blue) and 100% glycated/100% crosslinked (red) protein. Solid lines are the average of 10 independent simulations. Shaded areas represent SD. **(N)** Heatmap representing N2BA titin length at 10 pN peak force for Monte Carlo simulations at different glycation conditions, relative to the non-modified protein. Results are the average of 10 independent simulations per condition. The simulations in panels M and N consider that the PEVK region can be glycated. Error bars are SD of bootstrapping distributions^75^. PDF: probability density function.

### Mechanical properties of glycated titin domains

Although intramolecular crosslinks are generally considered to stiffen proteins due to the associated reduction in total contour length, this effect can be countered or even reversed depending on how they affect the mechanical (un)folding dynamics of the targeted domains; for instance if crosslinks result in mechanical destabilization that favors unfolded polypeptides^43^. Hence, to quantify in detail how crosslinking AGEs affect the global mechanical properties of titin domains, as well as the contribution of non-crosslinking AGEs, we further exploited our single-molecule force-ramp experiments.

First, we determined whether AGEs alter the mechanical stability of the I91 domain. We find that domains containing MG-derived crosslinking AGEs unfold at a slightly lower force than counterparts incubated in the absence of MG (142 and 158 pN, respectively; **Figure 6C**). Similarly, CEL-containing I91 domains appear mechanically weaker than the corresponding controls (114 vs 122 pN, respectively; **Figure 6D**). Fitting the Bell-Evans model of force-activated reactions^55^, we estimate that unfolding rates at zero force are between 3 and 8 fold higher for glycated domains, while the distance to the transition state is mostly preserved indicating no major changes in the force dependency of unfolding (**Table S2**). In combination, our analyses of unfolding forces indicate that both crosslinking and non-crosslinking AGEs induce mechanical weakening of the targeted domains, a softening effect that in the case of crosslinking modifications opposes stiffening that results from reduced total contour length.

Next, we did unfolding-quench-probe AFS experiments to study mechanical folding of glycated I91 (**Figure 6E**)^29^. In these experiments, (I91)_8_ is subjected to an unfolding force-ramp, followed by a quench pulse to 0 pN where domains first collapse and subsequently regain native mechanical stability in a time-dependent manner. In the final probe pulse, domains that refolded during the quench pulse unfold again. Folding fractions are calculated as the ratio between the number of unfolding events observed in the probe pulse and those observed in the unfolding pulse. To have better resolution of folding fractions at low folding times, we included force pulses to 40 pN before and after the quench at 0 pN. Since 40 pN is non-permissive for folding of the I91 domain^41^, the initial pulse allows better synchronization of actual folding times limiting the effects of variable collapse times. Similarly, jumping straight to 40 pN avoids folding reactions occurring during the probe force ramp. Results of the unfolding-quench-probe AFS experiments indicate that CEL moieties do not noticeably modify the folding kinetics of the I91 domain (**Figure 6F**). In contrast, I91 domains containing crosslinking AGEs refold at a rate ≥30 s^-1^, which is the resolution limit of our setup implying at least a 6-fold increase compared to the corresponding control (**Figure 6G**). Hence, in addition to stiffening resulting from reduced total contour length, crosslinking AGEs entail an additional stiffening effect as a consequence of increased folding of targeted domains.

### Prevailing stiffening induced by titin glycation

As explained in the previous section, our single-molecule data demonstrate that, similar to oxidative modifications, glycation of titin can entail both stiffening and softening effects^29,42^. To illustrate the available range of modulation of titin mechanics by AGEs, we did Monte Carlo computer simulations as reported^43^. We first built virtual models of the I-band of human titin containing 104 (for the N2BA isoform) or 48 (for the N2B isoform) Ig domains, all of them with equivalent mechanical properties, as well as random coil N2Bus and PEVK regions (**Table S3**). Since the PEVK region is rich in lysine residues (**Figure 6H,I**), we also estimated the reduction in contour length associated with crosslinking modifications targeting this region of titin. Specifically, considering that the first and last lysine in each of the 31 PEVK repetitions can form a crosslink (**Figure 6I**), we used a 67% maximum reduction in contour length upon glycation of the PEVK region (**Figure 6J**). We acknowledge that crosslinks between different PEVK repetitions could result in further decreases in contour length; however, we have not contemplated this possibility here.

In the simulations, we subjected the virtual I-bands to 1-Hz triangular force pulses between 0 pN and a predefined setpoint peak force while monitoring the resulting changes in titin length (**Figure 6K,L**). During these extension/relaxation cycles, titin domains unfold and refold stochastically according to their folding and unfolding rates, which depend on the type of glycation modification as detected in our single-molecule experiments (**Tables S4, S5**). At t = 0 s, all domains are folded and, as the simulations progress, a fraction of domains transitions to the unfolded state resulting in longer protein lengths (**Figure 6L**). Simulation times were long enough to always reach steady-state lengths in all simulation runs (**Figures 6M, S13**).

In **Figure 6M**, we present the results of the simulations of N2BA titin at a peak force of 10 pN, typically considered the upper limit of the physiological range^40^, comparing control conditions with two extreme scenarios where all domains are glycated with either non-crosslinking or crosslinking AGEs. These results readily capture the prevailing stiffening effect of crosslinking modifications, which cause ∼33% shortening of titin at peak force. In contrast, non-crosslinking modifications result in subtle, yet measurable, softening. Prompted by these results, we ran additional simulations at intermediate degrees of glycation and considering different proportions of non-crosslinking and crosslinking AGEs. In **Figure 6N**, we compare steady-state lengths of titin in these simulations with respect to control simulations. As expected, we find that the mechanical effects of both types of AGEs progressively increase at higher glycation fractions. More importantly, we observe that stiffening contributions of crosslinking AGEs outweigh softening induced by non-crosslinking counterparts even when the ratio of crosslinking/non-crosslinking modifications is only 20/80, independently of the total levels of glycation (**Figure 6N**). We observe similar effects for the N2B isoform, although in this case stiffening is less apparent (**Figure S13 A,B**). Increasing the peak force of our simulations to 100 pN exacerbates the mechanical consequences of both crosslinking and non-crosslinking modifications, with up to 57% reduction in length induced by crosslinking AGEs (**Figure S13C-F**). Further analysis of the Monte Carlo simulations captures a fundamental role of glycation of the PEVK domains in global titin stiffening (**Figure S14**). Indeed, if we consider that no glycation targets the PEVK region, no stiffening of titin is detected at 10 pN peak force indicating no overall mechanical effect of crosslinking AGEs targeting Ig domains at this force (**Figure S14A,C**). At 100 pN in the absence of PEVK glycation, the stiffening contribution of crosslinking AGEs in Ig domains becomes apparent again (**Figure S14B,D**). Such force dependency of the stiffening effects of crosslinking AGEs in titin Ig domains resembles the behavior reported for intradomain disulfide bonds^43^.

Finally, we also quantified from our Monte Carlo simulations how glycation affects Ig domain unfolding/refolding dynamics, which can influence downstream force-dependent interactions and PTMs^11,15^ and active force generation by sarcomeres^41^. With this aim, we counted the number of unfolding events in the different simulations. Results indicate that both crosslinking and non-crosslinking AGEs increase the extent of Ig domain (un)folding transitions, particularly at low forces (**Figure S15**).

## Discussion

High concentrations of carbohydrates and the associated toxic side-products of glycolysis (e.g. α-oxoaldehydes) are at the basis of many human diseases including diabetes, obesity, age-related disorders and the metabolic syndrome, as well as their associated comorbidities (e.g. cardiovascular complications)^56^. However, the underlying molecular pathomechanisms are yet to be fully elucidated. A main example is the origin of myocardial stiffening typically associated with carbohydrate-related diseases, a major risk factor for developing heart failure^3^. While fibrosis and glycation of ECM components have been shown to stiffen the myocardium in these conditions^6,7,27^, the reasons for the concurrent rigidification of cardiomyocytes (even permeabilized ones) remain incompletely understood, limiting our grasp on how diastolic failure develops in affected patients. So far, stiffening of diabetic cardiomyocytes has been proposed to be contributed by hypophosphorylation of titin and by glycogen accumulation, although compensatory overexpression of the compliant N2BA titin isoform complicates quantitative interpretation^6,33,57,58^. The data presented here uncover an additional stiffening effect induced by titin glycation, a modification found in the myocardium of both animal models and humans in diabetes^34^ (**Figure 1C-D**) and aging^35^.

Specifically, our work demonstrates that MG-induced glycation, a main chemical route during glycative stress^20^, results in cardiomyocyte stiffening (**Figure 1E-I**) because it leads to the preferential formation of crosslinking AGEs within titin domains. Indeed, Monte Carlo simulations indicate that stiffening induced by intradomain crosslinking AGEs prevails over softening contributions by non-crosslinking counterparts already when the proportion of crosslink-containing domains reaches 20% (for N2BA titin) or 40% (for N2B titin), independently of the total extent of glycation (**Figures 6N, S13**). From our single-molecule experiments at saturating glycation conditions, we estimate the actual proportion of crosslink-containing domains to be at least close to 80% (**Figures 2N, S4**), which agrees with the observation using WB that the percentage of CEL within all the MG-derived AGEs is relatively low (**Figure 4C**). This level of crosslinking modifications consistently leads to titin stiffening in Monte Carlo simulations, in line with our cell mechanics experiments. Since simulations also show that glycation of the PEVK region is a main contributor to titin stiffening (**Figure S14**), the different proportions of crosslink-containing domains required to achieve stiffening in N2BA and N2B titin can be easily explained by the distinct relative mechanical contribution of the PEVK domains in both isoforms (the ratio PEVK length in nm/number of Ig domains is 6.9 and 1.4 for the N2BA and the N2B isoforms, respectively) (**Table S3**). To strengthen our conclusions, we have obtained further evidence of the presence of crosslinking AGEs in MG-treated titin domains by mass spectrometry (**Figure 5**) and NMR (**Figure 3C**). Furthermore, our NMR results show that the structure of I91 is mostly unaffected by MG-induced glycation (**Figures 3D, S7**), which induces instead restricted conformational entropy of the protein as a result of the formation of crosslinking AGEs (**Figures 3F-H, S8A, S9**). Such overall preservation of the I91 native fold is also supported by the modest reduction in mechanical stability of glycated domains (**Figure 6C**) and their enhanced/preserved ability to refold (**Figure 6F,G**). Hence, in combination, our findings add to available evidence challenging the conventional view that glycation universally compromises protein structure^17^.

A potential limitation of our work stems from the millimolar MG concentration we used to trigger glycation in timescales compatible with single-molecule and cell mechanics experiments. This concentration is several orders of magnitude above typical estimates in cells, which indicate that MG levels do not exceed the micromolar range even in situations of glycative stress^59^. Hence, it appears that the increased glycation of titin in diabetes and aging that we and others have observed (**Figure 1C,D**)^34,35^ most probably results from the slow accumulation of AGEs involving multiple glycation pathways, a situation challenging to reproduce in the laboratory. Still, we have been able to detect a measurable fraction of crosslinking AGEs occurring very rapidly after addition of 50 mM MG (**Figure 2N**). This observation points to favored glycation reactions induced by MG that can proceed at physiological concentrations. It is also important to consider that given the high reactivity of MG and that the glycolytic flux can be estimated to produce ∼125 μmol of MG per kg of cell mass per day (i.e. 133 μM per day for a 15,000 μm^3^ cardiomyocyte considering a cellular density of 1.06 g/mL)^20^, it is conceivable that the effects of any given steady-state concentration of MG *in vivo* are noticeably higher than those of a seemingly equivalent, but not replenishable, concentration of MG in *in vitro* experiments.

Several aspects of our work call for further research. First, given that intracellular protein glycation is common to many tissues in addition to the myocardium^60^, we speculate that equivalent mechanisms to the ones we describe here can contribute to stiffening of cell types other than cardiomyocytes. Similarly, it is possible that glycation targeting protein-folding-based mechanosensor proteins like talin can have profound consequences in cell behavior^61^. In this regard, although the existence of crosslinking AGEs can be extrapolated from the presence of non-crosslinking counterparts^19^, new methods are required to quantify crosslinking modifications in native proteins and how they depend on cellular state. These new methods will need to overcome both the vast combinatorial space of potential target sites, particularly in large proteins like titin, and the interpretation and scoring of hybrid fragmentation spectra, which is very challenging for conventional mass-spectrometry approaches^62,63^. For further quantitative understanding of the effects of protein AGEs in cell mechanics, additional work will need to address protein folding intermediates^41^ and to integrate both stiffening interdomain crosslinks and the effects of mechanical crosstalk with other PTMs, especially those competing for glycation sites like acetylation and ubiquitination^64^.

## Concluding remarks

Our results with titin add to the list of detrimental functional consequences that follow glycation of key proteins in cardiomyocytes^64,65^, including other sarcomere components important for contraction^34,66–68^, the calcium pump SERCA2a^69^ and the calcium channel RyR2^35^. At the methodological level, our work illustrates how single-molecule AFS can discriminate the mechanical consequences of crosslinking and non-crosslinking AGEs in a manner that is chemically agnostic, therefore overcoming intrinsic heterogeneity of non-enzymatic glycation reactions. Having pinpointed crosslinking modifications as culprits of cell rigidification, our work provides deeper understanding of highly promising therapeutic strategies based on scavenger molecules that block protein glycation^27^. According to our data, further improvements could be achieved by preferentially preventing the formation of crosslinking AGEs, or even by inducing their cleavage. Single-molecule force spectroscopy methods can serve as a valuable screening platform to this aim.

## Author contributions

Conceptualization: A.B., A.C.D., E.H.G., M.A., Jor.A.C.

Methodology: A.C.M., E.C., R.S.R., N.V., D.S.O., I.M.M., D.V.C., M.R.P., F.M.E., Joa.A.C., C.P.M., R.G., J.V., I.F.P.

Investigation: A.B., A.C.D., A.C.M., E.C., E.H.G., M.A., Jor.A.C.

Funding acquisition: J.V., Jor.A.C.

Writing – original draft: A.B., A.C.D., M.A., Jor.A.C.

Writing – review & editing: all authors

## Competing interests

Authors declare that they have no competing interests.

## Data availability

The authors declare that the data supporting the findings of this study are available within the paper and its Supplementary Information files. Should any raw data files be needed in another format they are available from the corresponding authors upon reasonable request. The NMR assignments have been deposited in the Biological Magnetic Resonance Data Bank (BMRB) under the accession codes 53389 and 53390 for (I91)_1_ and MG-treated (I91)_1_, respectively.

## Code availability

The custom-built code used in this study is available from the corresponding authors upon reasonable request.

## Acknowledgements

Jor.A.C. acknowledges funding from Ministerio de Ciencia, Innovación y Universidades (MICIU, MICIU/AEI/10.13039/501100011033) through grants PID2020-120426GB-I00, PID2023-147683NB-I00, PLEC2022-009235 and RED2022-134242-T, the Regional Government of Madrid (grant Tec4Bio S2018/NMT-4443, 50% co-financed by the European Social Fund and the European Regional Development Fund for the programming period 2014-2020, and grant TecNanoBio TEC-2024/TEC-158, *Bases Reguladoras* 2402/2024; Convocatoria 3177/2024), and the European Research Council (ERC) under the European Union’s Horizon 2020 research and innovation program (grant agreement No. [101002927]). J.V. acknowledges grants PID2021-122348NB-I00 and PID2021-126827OB-I00 funded by MICIU and by “ERDF A way of making Europe”, PLEC2022-009298, PLEC2022-009235 and EQC2021-007053-P funded by MICIU and by “European Union NextGenerationEU/ PRTR”, and S2022/BMD-7333-CM (INMUNOVAR-CM) funded by Comunidad de Madrid. The project leading to these results has received funding from ‘la Caixa’ Foundation under the project code LCF/PR/HR22/52420019. CNIC is supported by the Instituto de Salud Carlos III (ISCIII), the MICIU and the Pro CNIC Foundation, and is a Severo Ochoa Center of Excellence (grant CEX2020-001041-S funded by MICIU). A.B. was the recipient of an FPI predoctoral fellowship (PRE2018-084392 funded by MICIU). A.C.D. acknowledges funding from ‘la Caixa’ Foundation (LCF/BQ/PI22/11910029) and the MICIU through project PID2022-140352NA-I00. A.C.D. is a Research Associate of the Belgian FNRS. A.C.M. is a recipient of a predoctoral fellowship funded by the Regional Government of Madrid (PIPF-2023SAL-GL-31131). R.S.R. acknowledges funding from the European Molecular Biology Organization (EMBO, postdoctoral fellowship EMBO ALTF 417-2022). I.M.M. was the recipient of a Doctoral INPhINIT fellowship from ‘la Caixa’ Foundation (ID100010434, fellowship code LCF/BQ/DR20/11790009). I.F.P. acknowledges funding from Fundação para a Ciência e Tecnologia (2023.17125.ICDT). We thank the personnel from CNIC animal housing, Matthew M. Borkowski (Aurora Scientific) for his help with the Permeabilized Myocyte Test System 1600A, Guadalupe Sabio and María Villalba for their advice on animal models, Jaime Andrés Rivas-Pardo for advice on titin purification and Jonathan A. Kirk and Álvaro Martínez-del-Pozo for feedback. We thank Gabriel Martorell and Rosa Gomila, both from the Scientific and Technical Services of the University of the Balearic Islands, for their assistance in performing NMR and MALDI-TOF/TOF experiments. We thank all members of the Molecular Mechanics of the Cardiovascular System team for their support and input. The authors thank the donors and the Hospital Universitario Puerta de Hierro Majadahonda (HUPHM)/Instituto de Investigación Sanitaria Puerta de Hierro-Segovia de Arana (IDIPHISA) Biobank (Carlos III Health Institute Biomodels and Biobanks Platform – PT23/00015) for the human specimens used in this study.

## Materials and methods

### Human subject research

Samples and data from patients included in this study (**Table S6**) were provided by the Hospital Universitario Puerta de Hierro Majadahonda (HUPHM)/Instituto de Investigación Sanitaria Puerta de Hierro-Segovia de Arana (IDIPHISA) Biobank (Carlos III Health Institute Biomodels and Biobanks Platform – PT23/00015). Samples were processed following standard operating procedures with the appropriate approval of the Ethics and Scientific Committee (Ref: 67/187906.9/24). The handling of samples and patient data was conducted in accordance with the principles of the Declaration of Helsinki and the International Conference on Harmonization of Good Clinical Practice guidelines, under the framework of Spanish and European regulations concerning data protection and clinical research.

### Animal research

Mice were housed and maintained in CNIC’s animal facility in accordance with Spanish and European Legislation (Directive 2010/63/EU amended by Regulation EU 2019/1010; CNIC-01/18-PROEX042/1; CNIC-01/23 - PROEX 107.8/23). Only male mice were used for the experiments. Adult mice were sacrificed using CO_2_, and neonates were sacrificed by decapitation. Hearts were isolated, perfused with PBS, frozen in liquid nitrogen, and stored at - 80°C. All procedures on rats were reviewed and approved by the Faculty of Medicine of the University of Porto (FMUP) Animal Welfare and Ethics Review Body (Órgão Responsável pelo Bem-Estar dos Animais [ORBEA-FMUP]) and the Portuguese competent authority (Direção Geral de Alimentação e Veterinária [DGAV], reference number 8161/23-S) and performed in accordance with EU Directive 2010/63/EU and Decreto-lei 113/2013 national legislation at the FMUP animal facility. Seven-week-old male Wistar-Han rats were acquired from Charles River Laboratories (Barcelona, Spain) and housed in groups of 4 animals per cage in a controlled environment under a 12:12-h light-dark cycle at a RT of 22°C, with free supply of food and water. Twelve weeks later, animals were anesthetized with sevoflurane (8%), the heart was removed and immediately frozen in liquid nitrogen for subsequent analysis. All procedures were carried out by properly trained and licensed researchers.

### LC-MS/MS mass spectrometry in native titin experiments

Myocardial protein extracts in the presence of N-ethylmaleimide were obtained following Herrero-Galán et al.^43^. Samples were run in 3.5% SDS-PAGE gels in the absence of reducing agents. Proteins were visualized with Coomassie blue and titin bands were sliced out from the gel and stored at 4°C until analysis. Bands were equilibrated in 50 mM ammonium bicarbonate (ABC) prior to reduction with 50 mM DTT and alkylation with 100 mM iodoacetamide, both in 100 mM ABC. Gel bands were then subjected to in-gel-digestion using modified bovine chymotrypsin, sequencing grade (Promega) at a final ratio of 1:20 (chymotrypsin-protein). Digestion proceeded overnight at 37°C in 100 mM ABC, pH 7.8. After digestion, peptides were extracted using acetonitrile with 0.1% (v/v) trifluoroacetic acid (TFA). Finally, peptides were desalted and dried until LC-MS analysis. LC-MS analysis was done using an Evosep One HPLC (Evosep) coupled to an Orbitrap Eclipse Tribrid Mass Spectrometer (Thermo Fisher Scientific) using an Endurance Evosep column 15 cm x 150 μm ID as analytical column (Thermo Fisher Scientific) coupled to a stainless steel emitter of 30 μm ID. Peptides were eluted from Evotips and analyzed using the Evosep One pre-programmed gradient for 15 samples per day (SPD). MS spectra were acquired in the Orbitrap analyzer using full ion-scan mode with a 390-1700 m/z range and 60,000 FT resolution. The automatic gain control target was set at 1 x 10^6^ with 50 ms maximum injection time. HCD fragmentation was performed at 30% of normalized collision energy and MS/MS spectra were analyzed at a 30,000 resolution in the Orbitrap with automatic gain control target set at 1 x 10^5^ and 100 ms maximum injection time. For peptide identification, tandem mass spectra were extracted, and charge state was deconvoluted by Proteome Discoverer 2.5.0.400 (Thermo Fisher Scientific). MS/MS spectra were analyzed using SEQUEST HT (Thermo Fisher Scientific) performing searches against a titin database^43^ considering as variable modifications methionine oxidation (Δmass = 15.995), NEM and IAM-modified cysteines (Δmass = 125.048 and 57.021, respectively), and the masses of non-crosslinking AGEs CML (Δmass = 58.005), CEL (Δmass = 72.021) and MG-H (Δmass = 54.011). Peptide-spectrum matches (PSM) were filtered to a q-value < 0.01.

### Purification of methylglyoxal

MG was purchased as a 40% solution (Sigma-Aldrich) and additionally purified by steam distillation. The concentration of MG after purification was quantified by incubating with excess H_2_O_2_ and titrating the remaining H_2_O_2_ with KMnO_4_ according to Friedemann’s method^70^. The fractions were frozen until use.

### AFM-based nanomechanical spectroscopy on neonatal cardiomyocytes

Dissected hearts from neonatal (postnatal days 1-3) mice were minced with scissors in cold Hanks’ Balanced Salt Solution (HBSS) and dissociated using the Pierce Primary Cardiomyocyte Isolation Kit (Thermo Fisher Scientific) in 0.21 mL enzyme mix/heart in a 2 mL reaction tube for 20 min at 37°C under constant end-to-end rotation. The pellet was centrifuged, washed twice with HBSS and resuspended in 0.5 mL cardiomyocyte medium (DMEM for Primary Cell Isolation containing 10% heat-inactivated FBS and 1% penicillin/streptomycin)/heart and plated onto MatTek dishes covered with matrigel solution (Fisher Scientific). AFM measurements were performed with a commercial JPK NanoWizard III instrument (Bruker-JPK) coupled to an inverted Axio Observer A1 optical microscope (Carl Zeiss). Cardiomyocytes were permeabilized with 0.2% (v/v) Triton X-100 in 1% (w/v) BSA for 15 min and incubated with 50 mM MG or PBS for 4 h at 37°C. Following rinsing with PBS, cardiomyocytes were probed at RT in PBS using rectangular Si_3_N_4_ AFM cantilevers with silicon tips of radius < 15 nm and a nominal spring constant of 0.1 N m^-1^ (BioLever mini, Bruker). Cantilevers were calibrated using the thermal fluctuation method^71^. FD curves were recorded in contact mode on the surface of single cardiomyocytes to determine Young’s moduli. Approach-retract cycles were performed with a tip velocity of 10 µm s^-1^, a ramp size of 2 µm and a setpoint force of 2 nN. We analyzed around 100 FD curves per cardiomyocyte. Young’s modulus values (*E*) were calculated using the JPK Data Analysis software, by fitting the approach section of the curve to the Hertz model using Equation _1_^72^:

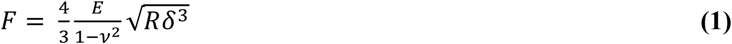

In Equation 1, *F* is the applied force, *R* is the probe radius, *ν* is Poisson’s ratio, and *δ* is the indentation. Cells mostly consist of water and are considered incompressible, hence a Poisson ratio of 0.5 was used.

### Tensile testing of adult rat cardiomyocytes

Rat cardiomyocytes were isolated from cardiac tissue of 19-week-old male Wistar rats. Cells were isolated in imidazole-containing relaxing buffer (RB, **Table S7**) by homogenizing a piece of tissue from the left ventricle with ceramic microspheres (Bertin Instruments) for 10 seconds at 5,000 rpm using a MiniLys system (Bertin Instruments). The cell suspension was resuspended to a final volume of 2.5 mL. Cells were incubated with 0.5% Triton X-100 (Millipore) for 5 min on ice. Subsequently, the detergent was removed by several washes in 15 mL imidazole-containing RB followed by centrifugation of the cells for 1 min at 348 *g*. Passive tension measurements were performed using a Permeabilized Myocyte Test System 1600A (Aurora Scientific), where single isolated skinned cardiomyocytes were fixed to a force transducer and a motor. Cells were transferred to propionic-containing RB (**Table S7**) and passive force was measured during 6 step length increments of 5% L_0_. To avoid quenching of MG with phosphocreatine, cells were incubated with MG (or controls in the absence of MG) for 30 min at in phosphocreatine-free RB (**Table S7**). Finally, the cells were transferred back to propionic-containing RB and the passive tension was measured again. All force values were converted to tension values assuming an elliptical shape of the cell, according to

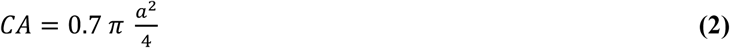

where *a* is the width of the cardiomyocyte observed in the microscope and CA is the estimated cross-sectional area^73^. In control experiments to check for the potential effect of ATP modification by MG, passive tension of cardiomyocytes was first measured in phosphocreatine-free RB. Next, cells were incubated for 30 min in a phosphocreatine-free RB that had been previously incubated for 30 min with 50 mM MG (excess MG in the buffer was quenched with 0.1 M Tris-HCl pH 7.5; *ATP-modified RB*), and passive tension was determined. To account for non-specific effects of adducts between Tris and MG, we included a control experiment where Tris was added to MG before final addition to phosphocreatine-free RB (*ATP-preserved RB*).

### Protein expression and purification

The complementary DNA (cDNA) coding for monomers and octamers of titin I91 was cloned in the pQE80 expression plasmid (Qiagen). The cDNA coding for titin (I91-K6/37/79A)_8_ synthetized by GeneArt Gene Synthesis (Thermo Fisher Scientific) was inserted in the pQE80 expression plasmid using BamHI and KpnI restriction enzymes and verified by Sanger sequencing. Polyproteins were expressed in *Escherichia coli* BLR (DE3). Monomeric protein was expressed in *Escherichia coli* BL21 (DE3) and cultured in isotopically labeled M9 minimal medium (**Table S8**) containing ^13^C_6_-D-glucose and ^15^N-ammonium chloride. Fresh cultures (OD_600_ = 0.6-1.0) were induced with 1 mM isopropyl β-D-1-thiogalactopyranoside (IPTG) for 3 h at 37°C and 250 rpm agitation. Bacteria culture was lysed using French Press. Proteins were purified by affinity chromatography using gravity-based strategy or in a HisTrap Fast Flow column using Fast Protein Liquid Chromatography (FPLC) AKTA Pure 25 L system (GE Healthcare). Proteins were eluted in 50 mM sodium phosphate pH 7, 300 mM NaCl, 250 mM imidazole, followed by protein concentration using Amicon Ultracel 3K, for (I91)_1_, or 30K, for (I91)_8_, centrifugal filter (Millipore). Finally, we did an additional purification step using size-exclusion chromatography in Superdex 200 Increase 10/300 GL or Superose 6 Increase 10/300 GL columns in the AKTA Pure 25 L system in Hepes (10 mM Hepes, 150 mM NaCl, 1 mM EDTA, pH 7.2) or in 0.2 M phosphate (pH 7.4). Purity of samples was checked by SDS-PAGE. Protein concentration determinations considered theoretical extinction coefficients values (**Table S9**).

### Protein glycation

For batch glycation, Ni-NTA fractions with the highest protein concentration were pulled together, followed by buffer exchange to 0.2 M phosphate pH 7.4 using PD-10 desalting columns (GE Healthcare) or by dialysis using Slide-A-Lyzer Dialysis Casettes (Thermo Scientfic) with 3,500 Da molecular weight cut-off (MWCO) for (I91)_1_ and 20,000 Da MWCO for (I91)_8_ proteins for 24 h at 4°C, 100 rpm agitation. Reactions with MG were done at 37°C and excess MG was removed by size-exclusion chromatography using FPLC as described above. To induce formation of CEL, the protein was incubated with 70 mM pyruvic acid (Sigma-Aldrich) and 100 mM sodium cyanoborohydrate (NaBH_3_CN) (ThermoFisher Scientific) for 48 h at 50°C ^49^, and excess reagents were removed as above. For proteins treated both with pyruvic acid and MG, a dialysis step in 0.2 M phosphate buffer pH 7.4 was included between both reaction steps.

### MALDI-TOF/TOF mass spectrometry

Protein preparations were dialyzed against mili-Q water. For (I91)_1_, 2 μl of the protein solution were mixed in a 1:1 ratio with a matrix solution containing 5 mg/mL of α-cyano-4-hydroxycinnamic acid prepared in a water:acetonitrile (70:30). For (I91)_8_, the protein solution was mixed in a 1:1 ratio with a matrix solution containing 10 mg/mL of sinapinic acid prepared in a water:acetonitrile (50:50). Both matrix solutions contained 0.1% TFA. 0.5 μL aliquots of those mixtures were spotted onto a steel target plate (MTP 384), air-dried, and subjected to mass determination. Mass spectra were acquired on a Bruker Autoflex III MALDI-TOF/TOF spectrometer equipped with a 200-Hz smart-beam pulsed N_2_ laser (λ 337 nm). The IS1 and IS2 voltages were 20 kV and 18.40 kV respectively, and the lens voltage was 8.4 kV. Measurements were performed using a positive reflector mode with matrix suppression below 4000 Da. External calibration was performed using a standard protein mixture.

### Single-molecule force spectroscopy by Atomic Force Microscopy

Single-molecule AFS measurements were done in a Luigs & Neumann setup following published protocols^44^. Briefly, 5-10 μL of purified protein solution were spread on 15-mm-diameter coverslips coated with 100-nm-thick gold (Luigs & Neumann). Experiments were performed in 0.2 M phosphate buffer pH 7.4 using silicon nitride MLCT-FB cantilevers (Bruker), coated with 60 nm titanium-gold on their back side. Cantilevers were calibrated by the thermal noise method^71^. Single polyproteins were picked up by pressing the tip against the coverslip for 1 second at 1,500 pN contact force. To trigger protein unfolding, a 40 pN s^-1^ force ramp for 6.5 seconds was applied. For data selection and analysis, two levels of fingerprinting were used to select single-molecule events. As a non-stringent fingerprint, all the traces containing at least one step of ∼25 nm, which corresponds to one I91 unfolding event^29^, were selected. The step size of all events appearing on the selected traces were measured and plotted on bidimensional histograms. Based on the results, a more stringent fingerprint criterion, by which only traces containing at least 2 events of 23-27 nm and/or 9-14 nm steps were selected, was used for further analysis. For analysis of mechanical unfolding, only traces presenting detachment at forces higher than 240 pN were selected to ensure full unfolding of the polyprotein. Traces presenting nonspecific events at forces >50 pN were discarded. Traces with more than 8 events were only considered for analysis in **Figure S12C**. To quantify the fraction of glycated protein domains, we divided the number of short steps by the total number of steps. For quantifying mechanical stability, we fit the AFS data to the Bell-Evans model, which considers that the mechanical unfolding rate of the protein (*r*) depends on the applied force (*F*) according to:

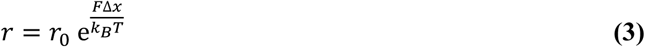

where *r*_0_ indicates the unfolding rate at zero force, Δ*x* is the distance to the transition state of the mechanical unfolding reaction, *k*_B_ is the Boltzmann constant and *T* is the absolute temperature^74^. Specifically, we used the following equation, which is the derivation of the Bell-Evans model for the case of a force ramp^55^:

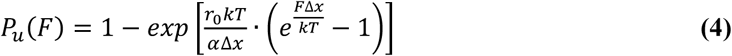

For folding analysis, a three-pulse experiment was used. Upon unfolding of domains in a 40 pN s^-1^ ramp, the force was quenched to 0 pN to allow for protein collapse and refolding. Finally, proteins were stretched again to detect domains that refolded during the quench time^29^. Folding fractions were calculated as the ratio between the number of unfolded domains during the probe pulse and the number of unfolded domains during the unfolding pulse. For this analysis, we only considered traces in which the difference in protein length between first and second ramp was not bigger than 15 nm. The folding ratio was quantified for varying quench times to quantify the folding rate. Errors were estimated using bootstrapping^75^.

### NMR spectroscopy

To prepare samples for NMR spectroscopy, ^13^C,^15^N-(I91)_1_ was incubated in the presence (*glycated sample*) or absence (*non-glycated sample*) of 50 mM MG for 48 h at 37°C. Solutions containing ^13^C,^15^N-(I91)_1_ (270 μM) or MG-treated ^13^C,^15^N-(I91)_1_ (200 μM) were prepared in 20 mM sodium acetate, pH 4.5 supplemented with 10% (v/v) D_2_O. In addition, the same buffer was used to prepare a solution containing non-labeled (I91)_1_ (120 μM) glycated with ^13^C-MG using the same conditions as above, and a solution containing 6 mM 1,3-bis((S)-5-amino-5-carboxypentyl)-5-methyl-1H-imidazol-3-ium acetic acid salt (sMOLD; Iris Biotech). These solutions were then used for NMR studies, which were carried out at 25°C on a Bruker Avance III spectrometer operating at a ^1^H resonance frequency of 600.1 MHz (14.1 T) and equipped with a 5 mm ^1^H/^13^C/D-BB z-GRD triple resonance broad band probe (TBI). In all experiments, water suppression was achieved by the watergate pulse sequence^76^ and proton chemical shifts were referenced to the water signal fixed at 4.771ppm. ^13^C and ^15^N chemical shifts were referenced indirectly using the ^1^H,X frequency ratios of the zero point^77^. The spectra were processed using the software packages NMRPipe/NMRDraw^78^ and Topspin (Bruker), whereas the data were analyzed using Xeasy/Cara^79^, Sparky^80^ and Protein Dynamics Center software (Bruker). The sequence-specific backbone assignments of native and MG-treated (I91)_1_, as well as the assignments of their C_β_, were achieved using the following standard 2D and 3D NMR experiments: ^1^H,^15^N-HSQC, ^15^N-TOCSY-HSQC, ^15^N-NOESY-HSQC (250 ms), HNCACB, CACB(CO)HN, HNCO, HCCH-TOCSY, HNHA, ^13^C-NOESY-HSQC and HN(CA)CO. To assign (I91)_1_ we initially started from part of the assignment previously published by Improta et al. at 35°C ^81^, which we adjusted, confirmed and expanded. The chemical shift assignment of MG-treated (I91)_1_ was achieved using the NMR experiments mentioned above. The NMR assignments have been deposited in the Biological Magnetic Resonance Data Bank (BMRB) under the accession codes 53389 and 53390 for (I91)_1_ and MG-treated (I91)_1_, respectively. Protein backbone assignments were used to estimate the secondary structure content at the residue level. This was carried out using three different algorithms: i) the neighbor corrected structure propensity calculator (ncSPC)^82^, which bases its calculation on the random coil library and adds an additional weighting procedure that accounts for the backbone conformational sensitivity of each amino acid type; ii) the TALOS+ program^83^, which uses the chemical shifts and the sequence information to make quantitative predictions of the secondary structural content; and iii) the CSI 3.0 web server, which uses backbone chemical shifts to identify up to eleven different types of secondary structures^84^. We also acquired ^15^N longitudinal (*R*_1_) and transverse (*R*_2_) relaxation data, as well as steady-state ^15^N HET-NOE data. The *R*_1_ values were determined using a series of 11 experiments with relaxation delays ranging from 10 to 2000 ms. The *R*_2_ values were obtained using a set of 11 different relaxation delays ranging from 2 to 140 ms. Recycle delays were 3 s in both *R*_1_ and *R*_2_ experiments. The ^15^N HET-NOE measurements were performed by 3 s high power pulse train saturation with a 5 s recycle delay. *R*_1_, *R*_2_ and ^15^N HET-NOE data were acquired using standard pulse sequences^85^. We acquired 32 scans in *R*_1_ and *R*_2_ experiments, and 200 scans in ^15^N HET-NOE experiment. *R*_1_ and *R*_2_ relaxation data were fitted to a mono-exponential decay function, whereas ^15^N HET-NOE data were obtained as the ratio of the peak intensities from the saturated and unsaturated spectra. Relaxation constants and experimental errors were calculated using the Protein Dynamics Center software (Bruker). *R*_1_ and *R*_2_ relaxation constants and their experimental errors were used to determine the correlation times (τ_c_) through the TENSOR2 software^86^. The orientation and the magnitude of the rotational diffusion tensor were determined from the *R*_1_ and *R*_2_ relaxation constants using the Quadratic-Diffusion software^87^. The experimental relaxation constants and the solution structure of I91 (PDB: 1TIT)^81^, which was modified adding the three last residues of our construct (i.e. R90-C92; **Text S1** using Pymol^88^), were studied using the molecular diffusion derived by Woessner in combination with the Lipari–Szabo model-free analysis of local flexibility^47^. The model-free order parameters (*O*^2^) report on the amplitudes of conformational fluctuations on time scales faster than the overall rotational diffusion. The amide bond length was fixed at 1.02 Å, whereas the chemical shift anisotropy was fixed at -172 ppm for the ^15^N backbone spins^89^. TENSOR2 software was also used to test five different models to characterize the internal dynamics of the amide groups: model 1 (*O*^2^), model 2 (*O*^2^, τ_e_), model 3 (*O*^2^, *k*_ex_), model 4 (*O*^2^, τ_e_, *k*_ex_) and model 5 (*O* ^2^, *O* ^2^, τ)^90^. τ is the effective internal correlation time (describes motions on a timescale > 20 ps); *k*_ex_ is a chemical exchange term (describes slow timescale motions on the order of μs–ms); and *O*_f_^2^ and *O* ^2^ are terms that result from splitting the generalized order parameter (*O*^2^) into two order parameters reflecting slower and faster motions, respectively. The confidence levels were estimated using 100 Monte Carlo simulations per run in combination with *c*^2^ and *F*-test criteria. The residues that had an overlapped cross-peak with other residues or they were too broadened to allow the quantitative analysis were excluded. The generalized order parameter (*O*^2^) was then used to estimate the change in the backbone conformational entropy (Δ*S*_conf_) of (I91)_1_ as a result of its modification with MG. This approach is based on the assumption that the fluctuations of individual residues are uncorrelated^91^, but it is also assumed that it underestimates the real change in the conformational entropy as it does not account for the entropy change associated with motional modes on time scales longer than τ_c_ (it only takes into account the *O*^2^ values). The estimation of the change in the backbone conformational entropy was carried out using

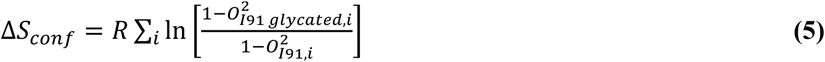

where *O*^2^ is the order parameter for each residue in native (I91)_1_ (*O*^2^_*I*91,*i*_) and in glycated (I91)_1_ (*O*^2^_*I*91,*glycated,i*_), and the sum runs over all residues^92^.

### NMR study of the formation of MOLD on I91

To assess the formation of the MG-derived crosslinking MOLD on MG-treated I91, we first assigned the ^1^H and the ^13^C-NMR chemical shifts of the imidazole ring of sMOLD (**Table S1**). This was achieved using a solution containing commercial MOLD (Iris Biotech) and collecting the corresponding 1D-^1^H, ^13^C-HSQC, ^13^C-edited-HSQC, ^1^H,^1^H-TOCSY, and ^13^C-HMBC spectra. Lastly, the obtained NMR data was used to identify the formation of MOLD on MG-treated I91, which was achieved collecting the ^13^C-HMBC spectrum of a solution containing non-labeled I91 that was previously incubated with ^13^C-MG (Sigma-Aldrich).

### Western blotting

Purified recombinant proteins were separated by 12% SDS-PAGE. Proteins were transferred onto polyvinylidene difluoride (PVDF) membranes (Biorad) and blocked in 5% non-fat dried milk solution in Tris-Buffered Saline with 0.1% Tween-20. Presence of glycation products was tested by 2 h incubation at RT with anti-CEL primary antibody (1:400, Cosmo Bio), followed by 1 h incubation at RT with HRP-conjugated anti-mouse secondary antibody (1:5000, ThermoFisher Scientific). Bands were visualized using ECL Western Blot Reagents (Cytiva).

### Analysis of recombinant (I91)_1_ by LC-MS/MS

Around 50 μg of protein were subjected to in-filter reduction and alkylation using iodoacetamide followed by chymotrypsin digestion (Nanosep Centrifugal Devices with Omega Membrane-30K, PALL), and the resulting peptides were desalted by C18 OMIX tips according to the manufacturer’s protocol, after which all fractions were dried under vacuum before MS analysis. MS analyses were performed as for native titin samples. To search for crosslinked lysines in the recombinant I91 titin sequence, we built a list of all possible MOLD-crosslinked combinations by proximity of lysine residues (K6-K55, K35-K79 and K37-K85) excluding combinations that would render the same mass as missed cleavages of non-modified peptides (**Table S10**). Different charge states for each combination were manually analyzed applying layouts with the QualBrowser 4.5.474.0 program (Thermo Xcalibur) by monitoring the retention time colocalization of 8 diagnostic fragments (according to the corresponding sequence), in the m/z window of the parental ion along the entire gradient. Scans containing the coeluting diagnostic-fragments for each combination were manually inspected to assign the remaining expected fragments resulting from the breaking of the crosslinked constructs, including the presence of signals belonging to expected neutral losses (ammonia loss, water loss and carbonyl-group loss generating b-derived “a” fragmentation series).

### Small-angle X-ray scattering

Solutions containing 140 μM (I91)_8_ that had been incubated in the presence or absence of MG were dialyzed against mili-Q water. Afterwards, these solutions were used for SAXS studies, which were carried out on a Xeuss 2.0 instrument (Xenocs) equipped with a microfocus Cu Kα source (λ 1.54 Å) and a Pilatus 300 k detector (Dectris). The distance between the detector and the sample was calibrated using silver behenate, and samples were measured using sample-detector distances of 370 mm. The measurements were carried out for 2 h under vacuum at 25°C using a BioCUBE device (Xenocs), equipped with a temperature-controlled flow-through quartz capillary (1.9 mm width). In addition, we also collected the scattering curve for milli-Q water, which was subtracted from the signal of the protein solutions. RAW software^93^ was used for data processing and analysis. GNOM was used to compute the pair-distance distribution functions (*P*r)^94^.

### Purification of titin from mouse cardiac tissue

Mouse cardiac tissue obtained by cryopulverization of frozen hearts in liquid nitrogen was used to extract titin following the protocol from^53^ with minor modifications. Buffer volumes were adjusted according to tissue weight (1mL per g of tissue). ∼ 200 mg of tissue were washed 4 times with 3 volumes of homogenization buffer (1 mM NaHCO_3_ pH 7, 50 mM KCl, 5 mM EGTA, 0.01 % NaN_3_) supplemented with protease inhibitors (1 mM PMSF, 0.1 mM leupeptin, 0.02 mM E-64, all from Sigma-Aldrich) and 20 µg/mL trypsin inhibitor (Roche). Between washes, fibers were pelleted by centrifugation (2,000 g, 10 min, 4°C). Then, fibers were resuspended in 2 volumes of extraction buffer (10 mM imidazole pH 7, 900 mM KCl, 2mM EGTA, 2 mM MgCl_2_, 0.01 % NaN_3_) supplemented with protease inhibitors (1.5 mM PMSF, 0.2 mM leupeptin, 0.04 mM E-64, 40 µg/mL trypsin inhibitor) and homogenized with a plastic microhomogenizer (Takara BioMasher) on ice. After homogenization, the tissue debris was pelleted by centrifugation (30 min, 20,000 g, 4°C) and the supernatant was diluted 4 times in precipitation buffer (0.1 mM NaHCO_3_ pH 7, 0.1 mM EGTA) supplemented with protease inhibitors (0.05 mM leupeptin, 0.01 mM E-64) and incubated for 1 h at 4°C in a spinning wheel. After incubation, samples were centrifuged (30 min, 20,000 g, 4°C) and the supernatant was diluted 5 times with precipitation buffer and incubated for 40 min at 4°C in a spinning wheel. Finally, the supernatants were centrifuged (40 min, 10,000 g, 4°C) to precipitate titin molecules and pellets were carefully resuspended in storage buffer (30 mM potassium phosphate pH 7.0, 200 mM KCl) at a ratio of 500 µl of buffer per 200 mg of cardiac tissue used as starting material. Resuspended titin molecules were stored at 4°C until use.

### Single-molecule force-spectroscopy on native titin

The experiments were carried out on HaloTag-ligand-coated surfaces, prepared according to published protocols^53^. Briefly, glass coverslips (15 mm, Ted Pella) were cleaned by sequential sonication (1% Hellmanex, acetone, 96% ethanol; 30 min each), air-dried, activated by exposure to O₂ plasma for 15 min, and silanized by immersion for 20 min in 1% (v/v) 3-aminopropyltrimethoxysilane (APTMS, Sigma-Aldrich) in ethanol. Unreacted silane was removed by ethanol rinses, then surfaces were air-dried, cured at 100°C for ≥1 h, and stored in a desiccator until use. For functionalization with HaloTag ligand, silanized coverslips were incubated overnight at RT in the dark with 1 mM HaloTag succinimidyl ester (O4, Promega) in 50 mM borax buffer (pH 8.5). After incubation, coverslips were washed with Milli-Q water, air-dried, and stored at 4°C in a humid chamber for up to one week. For titin unfolding experiments, 20 µL Halo–titin solution was applied to a HaloTag-ligand-coated surface, incubated for 10 min, and washed with phosphate buffer. Single-molecule AFM force spectroscopy was performed in force–extension mode in 0.2 M phosphate buffer pH 7.4 containing 50 mM MG or control buffer without MG, using MLCT Bio DC cantilevers (Bruker; nominal spring constant 0.01 N m⁻¹). Cantilevers were calibrated by the thermal noise method^71^. Single titin proteins were picked up by pressing the tip against the coverslip for 1 second at 2 nN contact force and unfolding was triggered by applying a ramp of 1.2 µm at a speed of 600 nm s⁻¹. Force–extension curves were baseline-corrected and peaks corresponding to titin domain unfolding were identified. The associated contour length increments (ΔLc) were estimated using the worm-like chain model^50^.

### Monte Carlo simulations

To estimate the effect of glycation on titin stiffness we applied Monte Carlo simulations to a model of titin’s I-band containing equivalent Ig domains (104 and 48 for the N2BA and N2B titin isoforms) and two entropic regions (PEVK and N2Bus)^43^. Associated contour lengths and Kuhn lengths are included in **Table S3**. To calculate the reduction in contour length by crosslinking modifications of the PEVK, we aligned the 31 domains of the PEVK region of human N2BA titin (Uniprot: Q8WZ42) using Clustal Omega and considered the possibility that the first and last lysine in each domain can form a crosslink. To calculate the associated reduction in contour length, we subtracted the positions of both lysines from the number of amino acid residues present in the domain, considering a contour length per amino acid of 0.4 nm^51^. Next, we added the contour length of MOLD, which we estimated to be 1.6 nm. As an example, for a PEVK domain that is 27 amino acid long and contains lysines at positions 6 and 27, the contour length upon crosslinking glycation would be (27-21) × 0.4 + 1.6 = 4 nm. Since the original contour length is 27 × 0.4 = 10.8 nm, the contour length is reduced by 63% in this example. For the simulations, we implemented an oscillating force protocol with triangular waves at a frequency of 1 Hz and 10 ms of simulation step for 1 h and measured the resulting length of titin. For each condition, we ran 10 independent simulations. Next, we averaged titin length at times longer than 300 s to ensure that steady state had been reached. Ratios between steady state lengths at different conditions were calculated and plotted as heatmaps. We used the Freely-Jointed Chain model^95^ to estimate lengths from contour length values. Unfolding events were considered as all-or-none events and their probability was calculated according to Bell’s model^74^. Parameters governing (un)folding kinetics of Ig domains are included in **Tables S4** and **S5**.

### Statistical analysis

The statistical analyses were conducted with either IGOR Pro or GraphPad Prism 8. The data are presented with the mean value either accompanied by the standard error of the mean (SEM) or standard deviation (SD), as specified. The statistical significance was determined by using Student T-tests. If experimental data did not follow a normal distribution, Mann-Whitney test was applied instead. Significance was defined as p-value <0.05 (*), <0.01(**), <0.001 (***).

## Notes

### Competing Interest Statement

The authors have declared no competing interest.

## References

1 Mammoto, T., Mammoto, A. & Ingber, D. E. Mechanobiology and developmental control. Annu Rev Cell Dev Biol 29, 27–61, (2013). <https://www.ncbi.nlm.nih.gov/pubmed/24099083>.

2 Paszek, M. J., Zahir, N., Johnson, K. R., Lakins, J. N., Rozenberg, G. I., Gefen, A., . . . Weaver, V. M. Tensional homeostasis and the malignant phenotype. Cancer Cell 8, 241–254, (2005). <https://www.ncbi.nlm.nih.gov/pubmed/16169468>.

3 Borlaug Barry, A., Sharma, K., Shah Sanjiv, J. & Ho Jennifer, E. Heart Failure With Preserved Ejection Fraction. JACC 81, 1810–1834, (2023). <10.1016/j.jacc.2023.01.049>.

4 Zuela-Sopilniak, N. & Lammerding, J. Can’t handle the stress? Mechanobiology and disease. Trends in Molecular Medicine 28, 710–725, (2022). <10.1016/j.molmed.2022.05.010>.

5 Lehman, S. J., Crocini, C. & Leinwand, L. A. Targeting the sarcomere in inherited cardiomyopathies. Nature Reviews Cardiology 19, 353–363, (2022). <10.1038/s41569-022-00682-0>.

6 Falcao-Pires, I., Hamdani, N., Borbely, A., Gavina, C., Schalkwijk, C. G., van der Velden, J., . . . Paulus, W. J. Diabetes mellitus worsens diastolic left ventricular dysfunction in aortic stenosis through altered myocardial structure and cardiomyocyte stiffness. Circulation 124, 1151–1159, (2011). <https://www.ncbi.nlm.nih.gov/pubmed/21844073>.

7 van Heerebeek, L., Hamdani, N., Handoko, M. L., Falcao-Pires, I., Musters, R. J., Kupreishvili, K., . . . Paulus, W. J. Diastolic stiffness of the failing diabetic heart: importance of fibrosis, advanced glycation end products, and myocyte resting tension. Circulation 117, 43–51, (2008). <https://www.ncbi.nlm.nih.gov/pubmed/18071071>.

8 Prenner, S. B. & Chirinos, J. A. Arterial stiffness in diabetes mellitus. Atherosclerosis 238, 370–379, (2015). <https://www.sciencedirect.com/science/article/pii/S0021915014016451>.

9 Loescher, C. M., Freundt, J. K., Unger, A., Hessel, A. L., Kuhn, M., Koser, F. & Linke, W. A. Titin governs myocardial passive stiffness with major support from microtubules and actin and the extracellular matrix. Nat Cardiovasc Res 2, 991–1002, (2023). <https://www.ncbi.nlm.nih.gov/pubmed/39196092>.

10 Bielajew, B. J., Hu, J. C. & Athanasiou, K. A. Collagen: quantification, biomechanics, and role of minor subtypes in cartilage. Nat Rev Mater 5, 730–747, (2020). <https://www.ncbi.nlm.nih.gov/pubmed/33996147>.

11 Linke, W. A. & Hamdani, N. Gigantic business: titin properties and function through thick and thin. Circ Res 114, 1052–1068, (2014). <https://www.ncbi.nlm.nih.gov/pubmed/24625729>.

12 Granzier, H. L. & Labeit, S. Discovery of Titin and Its Role in Heart Function and Disease. Circ Res 136, 135–157, (2025). <https://www.ncbi.nlm.nih.gov/pubmed/39745989>.

13 Martinez-Martin, I., Silva-Rojas, R. & Alegre-Cebollada, J. Mending the Achilles heels of titin in cardiac and musculoskeletal disease. Biophysical Reviews In press, (2026).

14 Alegre-Cebollada, J. Protein nanomechanics in biological context. Biophys Rev 13, 435–454, (2021). <https://www.ncbi.nlm.nih.gov/pubmed/34466164>.

15 Loescher, C. M., Breitkreuz, M., Li, Y., Nickel, A., Unger, A., Dietl, A., . . . Linke, W. A. Regulation of titin-based cardiac stiffness by unfolded domain oxidation (UnDOx). Proc Natl Acad Sci U S A 117, 24545–24556, (2020). <https://www.ncbi.nlm.nih.gov/pubmed/32929035>.

16 Binek, A., Castans, C., Jorge, I., Bagwan, N., Rodriguez, J. M., Fernandez-Jimenez, R., . . . Vazquez, J. Oxidative Post-translational Protein Modifications upon Ischemia/Reperfusion Injury. Antioxidants (Basel) 13, (2024). <https://www.ncbi.nlm.nih.gov/pubmed/38247530>.

17 Uceda, A. B., Marino, L., Casasnovas, R. & Adrover, M. An overview on glycation: molecular mechanisms, impact on proteins, pathogenesis, and inhibition. Biophys Rev 16, 189–218, (2024). <https://www.ncbi.nlm.nih.gov/pubmed/38737201>.

18 Fournet, M., Bonte, F. & Desmouliere, A. Glycation Damage: A Possible Hub for Major Pathophysiological Disorders and Aging. Aging Dis 9, 880–900, (2018). <https://www.ncbi.nlm.nih.gov/pubmed/30271665>.

19 Martin, M. S., Jacob-Dolan, J. W., Pham, V. T. T., Sjoblom, N. M. & Scheck, R. A. The chemical language of protein glycation. Nat Chem Biol 21, 324–336, (2025). <https://www.ncbi.nlm.nih.gov/pubmed/38942948>.

20 Schalkwijk, C. G. & Stehouwer, C. D. A. Methylglyoxal, a Highly Reactive Dicarbonyl Compound, in Diabetes, Its Vascular Complications, and Other Age-Related Diseases. Physiol Rev 100, 407–461, (2020). <https://www.ncbi.nlm.nih.gov/pubmed/31539311>.

21 Allaman, I., Belanger, M. & Magistretti, P. J. Methylglyoxal, the dark side of glycolysis. Front Neurosci 9, 23, (2015). <https://www.ncbi.nlm.nih.gov/pubmed/25709564>.

22 Thornalley, P. J. Dicarbonyl intermediates in the maillard reaction. Ann N Y Acad Sci 1043, 111–117, (2005). <https://www.ncbi.nlm.nih.gov/pubmed/16037229>.

23 Lai, S. W. T., Lopez Gonzalez, E. J., Zoukari, T., Ki, P. & Shuck, S. C. Methylglyoxal and Its Adducts: Induction, Repair, and Association with Disease. Chem Res Toxicol 35, 1720–1746, (2022). <https://www.ncbi.nlm.nih.gov/pubmed/36197742>.

24 Ahmed, M. U., Brinkmann Frye, E., Degenhardt, T. P., Thorpe, S. R. & Baynes, J. W. N-epsilon-(carboxyethyl)lysine, a product of the chemical modification of proteins by methylglyoxal, increases with age in human lens proteins. Biochem J 324 (Pt 2), 565–570, (1997). <https://www.ncbi.nlm.nih.gov/pubmed/9182719>.

25 Brinkmann, E., Wells-Knecht, K. J., Thorpe, S. R. & Baynes, J. W. Characterization of an imidazolium compound formed by reaction of methylglyoxal and Nα-hippuryllysine. Journal of the Chemical Society, Perkin Transactions 1, 2817–2818, (1995). <10.1039/P19950002817>.

26 Verzijl, N., DeGroot, J., Ben, Z. C., Brau-Benjamin, O., Maroudas, A., Bank, R. A., . . . TeKoppele, J. M. Crosslinking by advanced glycation end products increases the stiffness of the collagen network in human articular cartilage: a possible mechanism through which age is a risk factor for osteoarthritis. Arthritis Rheum 46, 114–123, (2002). <https://www.ncbi.nlm.nih.gov/pubmed/11822407>.

27 Norton, G. R., Candy, G. & Woodiwiss, A. J. Aminoguanidine Prevents the Decreased Myocardial Compliance Produced by Streptozotocin-Induced Diabetes Mellitus in Rats. Circulation 93, 1905–1912, (1996). <10.1161/01.CIR.93.10.1905>.

28 Sims, T. J., Rasmussen, L. M., Oxlund, H. & Bailey, A. J. The role of glycation cross-links in diabetic vascular stiffening. Diabetologia 39, 946–951, (1996). <https://www.ncbi.nlm.nih.gov/pubmed/8858217>.

29 Alegre-Cebollada, J., Kosuri, P., Giganti, D., Eckels, E., Rivas-Pardo, J. A., Hamdani, N., . . . Fernandez, J. M. S-glutathionylation of cryptic cysteines enhances titin elasticity by blocking protein folding. Cell 156, 1235–1246, (2014). <https://www.ncbi.nlm.nih.gov/pubmed/24630725>.

30 Targosz-Korecka, M., Jaglarz, M., Malek-Zietek, K. E., Gregorius, A., Zakrzewska, A., Sitek, B., . . . Szymonski, M. AFM-based detection of glycocalyx degradation and endothelial stiffening in the db/db mouse model of diabetes. Sci Rep 7, 15951, (2017). <https://www.ncbi.nlm.nih.gov/pubmed/29162916>.

31 Le Master, E., Fancher, I. S., Lee, J. & Levitan, I. Comparative analysis of endothelial cell and sub-endothelial cell elastic moduli in young and aged mice: Role of CD36. Journal of Biomechanics 76, 263–268, (2018). <https://www.sciencedirect.com/science/article/pii/S0021929018304366>.

32 Lieber, S. C., Aubry, N., Pain, J., Diaz, G., Kim, S. J. & Vatner, S. F. Aging increases stiffness of cardiac myocytes measured by atomic force microscopy nanoindentation. Am J Physiol Heart Circ Physiol 287, H645–651, (2004). <https://www.ncbi.nlm.nih.gov/pubmed/15044193>.

33 Hopf, A. E., Andresen, C., Kotter, S., Isic, M., Ulrich, K., Sahin, S., . . . Kruger, M. Diabetes-Induced Cardiomyocyte Passive Stiffening Is Caused by Impaired Insulin-Dependent Titin Modification and Can Be Modulated by Neuregulin-1. Circ Res 123, 342–355, (2018). <https://www.ncbi.nlm.nih.gov/pubmed/29760016>.

34 Papadaki, M., Holewinski, R. J., Previs, S. B., Martin, T. G., Stachowski, M. J., Li, A., . . . Kirk, J. A. Diabetes with heart failure increases methylglyoxal modifications in the sarcomere, which inhibit function. JCI Insight 3, (2018). <https://www.ncbi.nlm.nih.gov/pubmed/30333300>.

35 Ruiz-Meana, M., Minguet, M., Bou-Teen, D., Miro-Casas, E., Castans, C., Castellano, J., . . . Garcia-Dorado, D. Ryanodine Receptor Glycation Favors Mitochondrial Damage in the Senescent Heart. Circulation 139, 949–964, (2019). <https://www.ncbi.nlm.nih.gov/pubmed/30586718>.

36 Coleman, D. L. Diabetes-obesity syndromes in mice. Diabetes 31, 1–6, (1982). <https://www.ncbi.nlm.nih.gov/pubmed/7160533>.

37 Huxley, H. E. The mechanism of muscular contraction. Science 164, 1356–1365, (1969). <https://www.ncbi.nlm.nih.gov/pubmed/4181952>.

38 Han, S.-W., Kolb, J., Farman, G. P., Gohlke, J. & Granzier, H. L. Glycerol storage increases passive stiffness of muscle fibers through effects on titin extensibility. Journal of General Physiology 157, (2025). <10.1085/jgp.202413729>.

39 Wagner, F. A., Loescher, C. M., Unger, A., Kühn, M., Klotz, A. J., Liashkovich, I., . . . Linke, W. A. Direction-dependent contributions of cardiac myofilament networks to myocardial passive stiffness reveal a major disparity for titin. Basic Research in Cardiology, (2025). <10.1007/s00395-025-01119-8>.

40 Li, H., Linke, W. A., Oberhauser, A. F., Carrion-Vazquez, M., Kerkvliet, J. G., Lu, H., . . . Fernandez, J. M. Reverse engineering of the giant muscle protein titin. Nature 418, 998–1002, (2002). <https://www.ncbi.nlm.nih.gov/pubmed/12198551>.

41 Rivas-Pardo, J. A., Eckels, E. C., Popa, I., Kosuri, P., Linke, W. A. & Fernandez, J. M. Work Done by Titin Protein Folding Assists Muscle Contraction. Cell Rep 14, 1339–1347, (2016). <https://www.ncbi.nlm.nih.gov/pubmed/26854230>.

42 Giganti, D., Yan, K., Badilla, C. L., Fernandez, J. M. & Alegre-Cebollada, J. Disulfide isomerization reactions in titin immunoglobulin domains enable a mode of protein elasticity. Nat Commun 9, 185, (2018). <https://www.ncbi.nlm.nih.gov/pubmed/29330363>.

43 Herrero-Galan, E., Martinez-Martin, I., Sanchez-Gonzalez, C., Vicente, N., Bonzon-Kulichenko, E., Calvo, E., . . . Alegre-Cebollada, J. Basal oxidation of conserved cysteines modulates cardiac titin stiffness and dynamics. Redox Biol 52, 102306, (2022). <https://www.ncbi.nlm.nih.gov/pubmed/35367810>.

44 Popa, I., Kosuri, P., Alegre-Cebollada, J., Garcia-Manyes, S. & Fernandez, J. M. Force dependency of biochemical reactions measured by single-molecule force-clamp spectroscopy. Nat Protoc 8, 1261–1276, (2013). <https://www.ncbi.nlm.nih.gov/pubmed/23744288>.

45 Carl, P., Kwok, C. H., Manderson, G., Speicher, D. W. & Discher, D. E. Forced unfolding modulated by disulfide bonds in the Ig domains of a cell adhesion molecule. Proc Natl Acad Sci U S A 98, 1565–1570, (2001). <https://www.ncbi.nlm.nih.gov/pubmed/11171991>.

46 Alegre-Cebollada, J., Badilla, C. L. & Fernandez, J. M. Isopeptide bonds block the mechanical extension of pili in pathogenic Streptococcus pyogenes. J Biol Chem 285, 11235–11242, (2010). <https://www.ncbi.nlm.nih.gov/pubmed/20139067>.

47 Lipari, G. & Szabo, A. Model-free approach to the interpretation of nuclear magnetic resonance relaxation in macromolecules. 1. Theory and range of validity. Journal of the American Chemical Society 104, 4546–4559, (1982). <10.1021/ja00381a009>.

48 Li, Z., Raychaudhuri, S. & Wand, A. J. Insights into the local residual entropy of proteins provided by NMR relaxation. Protein Sci 5, 2647–2650, (1996). <https://www.ncbi.nlm.nih.gov/pubmed/8976574>.

49 Marino, L., Ramis, R., Casasnovas, R., Ortega-Castro, J., Vilanova, B., Frau, J. & Adrover, M. Unravelling the effect of N(epsilon)-(carboxyethyl)lysine on the conformation, dynamics and aggregation propensity of alpha-synuclein. Chem Sci 11, 3332–3344, (2020). <https://www.ncbi.nlm.nih.gov/pubmed/34122841>.

50 Bustamante, C., Marko, J. F., Siggia, E. D. & Smith, S. Entropic elasticity of lambda-phage DNA. Science 265, 1599–1600, (1994). <https://www.ncbi.nlm.nih.gov/pubmed/8079175>.

51 Ainavarapu, S. R., Brujic, J., Huang, H. H., Wiita, A. P., Lu, H., Li, L., . . . Fernandez, J. M. Contour length and refolding rate of a small protein controlled by engineered disulfide bonds. Biophys J 92, 225–233, (2007). <http://www.ncbi.nlm.nih.gov/pubmed/17028145>.

52 Rambo, R. P. & Tainer, J. A. Characterizing flexible and intrinsically unstructured biological macromolecules by SAS using the Porod-Debye law. Biopolymers 95, 559–571, (2011). <https://www.ncbi.nlm.nih.gov/pubmed/21509745>.

53 Rivas-Pardo, J. A., Li, Y., Mártonfalvi, Z., Tapia-Rojo, R., Unger, A., Fernández-Trasancos, Á., . . . Alegre-Cebollada, J. A HaloTag-TEV genetic cassette for mechanical phenotyping of proteins from tissues. Nature Communications 11, 2060, (2020). <10.1038/s41467-020-15465-9>.

54 Rief, M., Gautel, M., Oesterhelt, F., Fernandez, J. M. & Gaub, H. E. Reversible unfolding of individual titin immunoglobulin domains by AFM. Science 276, 1109–1112, (1997). <https://www.ncbi.nlm.nih.gov/pubmed/9148804>.

55 Schlierf, M., Li, H. & Fernandez, J. M. The unfolding kinetics of ubiquitin captured with single-molecule force-clamp techniques. Proc Natl Acad Sci U S A 101, 7299–7304, (2004). <https://www.ncbi.nlm.nih.gov/pubmed/15123816>.

56 Kroemer, G., Lopez-Otin, C., Madeo, F. & de Cabo, R. Carbotoxicity-Noxious Effects of Carbohydrates. Cell 175, 605–614, (2018). <https://www.ncbi.nlm.nih.gov/pubmed/30340032>.

57 Methawasin, M. & Granzier, H. Softening the Stressed Giant Titin in Diabetes Mellitus. Circ Res 123, 315–317, (2018). <https://www.ncbi.nlm.nih.gov/pubmed/30026374>.

58 Mellor, K. M., Varma, U., Koutsifeli, P., Curl, C. L., Janssens, J. V., Daniels, L. J., . . . Delbridge, L. M. D. Targeted glycophagy ATG8 therapy reverses diabetic heart disease in mice and in human engineered cardiac tissues. Nat Cardiovasc Res 4, 1487–1500, (2025). <https://www.ncbi.nlm.nih.gov/pubmed/41023280>.

59 Trujillo, M. N. & Galligan, J. J. Reconsidering the role of protein glycation in disease. Nature Chemical Biology 19, 922–927, (2023). <10.1038/s41589-023-01382-7>.

60 Di Sanzo, S., Spengler, K., Leheis, A., Kirkpatrick, J. M., Randler, T. L., Baldensperger, T., . . . Heller, R. Mapping protein carboxymethylation sites provides insights into their role in proteostasis and cell proliferation. Nat Commun 12, 6743, (2021). <https://www.ncbi.nlm.nih.gov/pubmed/34795246>.

61 Elosegui-Artola, A., Trepat, X. & Roca-Cusachs, P. Control of Mechanotransduction by Molecular Clutch Dynamics. Trends in Cell Biology 28, 356–367, (2018). <https://www.sciencedirect.com/science/article/pii/S0962892418300175>.

62 Leitner, A., Walzthoeni, T., Kahraman, A., Herzog, F., Rinner, O., Beck, M. & Aebersold, R. Probing native protein structures by chemical cross-linking, mass spectrometry, and bioinformatics. Mol Cell Proteomics 9, 1634–1649, (2010). <https://www.ncbi.nlm.nih.gov/pubmed/20360032>.

63 Trnka, M. J., Baker, P. R., Robinson, P. J., Burlingame, A. L. & Chalkley, R. J. Matching cross-linked peptide spectra: only as good as the worse identification. Mol Cell Proteomics 13, 420–434, (2014). <https://www.ncbi.nlm.nih.gov/pubmed/24335475>.

64 Delligatti, C. E. & Kirk, J. A. in Vitamins and Hormones Vol. 125 (ed Gerald Litwack) 47–88 (Academic Press, 2024).

65 LeWinter, M. M., Taatjes, D., Ashikaga, T., Palmer, B., Bishop, N., VanBuren, P., . . . Zile, M. Abundance, localization, and functional correlates of the advanced glycation end-product carboxymethyl lysine in human myocardium. Physiological Reports 5, e13462, (2017). <https://physoc.onlinelibrary.wiley.com/doi/abs/10.14814/phy2.13462>.

66 Janssens, J. V., Ma, B., Brimble, M. A., Van Eyk, J. E., Delbridge, L. M. D. & Mellor, K. M. Cardiac troponins may be irreversibly modified by glycation: novel potential mechanisms of cardiac performance modulation. Scientific Reports 8, 16084, (2018). <10.1038/s41598-018-33886-x>.

67 Papadaki, M., Kampaengsri, T., Barrick, S. K., Campbell, S. G., von Lewinski, D., Rainer, P. P., . . . Kirk, J. A. Myofilament glycation in diabetes reduces contractility by inhibiting tropomyosin movement, is rescued by cMyBPC domains. J Mol Cell Cardiol 162, 1–9, (2022). <https://www.ncbi.nlm.nih.gov/pubmed/34487755>.

68 Lewis, C. T. A., Moreno-Justicia, R., Savoure, L., Calvo, E., Bak, A., Laitila, J., . . . Ochala, J. Dysregulated skeletal muscle myosin super-relaxation and energetics in male participants with type 2 diabetes mellitus. Diabetologia 68, 1836–1850, (2025). <https://www.ncbi.nlm.nih.gov/pubmed/40295335>.

69 Shao, C. H., Capek, H. L., Patel, K. P., Wang, M., Tang, K., DeSouza, C., . . . Bidasee, K. R. Carbonylation Contributes to SERCA2a Activity Loss and Diastolic Dysfunction in a Rat Model of Type 1 Diabetes. Diabetes 60, 947–959, (2011). <10.2337/db10-1145>.

70 Friedemann, T. E. The action of alkali and hydrogen peroxide on glyoxals. Journal of Biological Chemistry 73, 331–334, (1927). <https://www.sciencedirect.com/science/article/pii/S0021925818843162>.

71 Hutter, J. L. & Bechhoefer, J. Calibration of atomic-force microscope tips. Review of Scientific Instruments 64, 1868–1873, (1993). <10.1063/1.1143970>.

72 Schillers, H., Rianna, C., Schape, J., Luque, T., Doschke, H., Walte, M., . . . Radmacher, M. Standardized Nanomechanical Atomic Force Microscopy Procedure (SNAP) for Measuring Soft and Biological Samples. Sci Rep 7, 5117, (2017). <https://www.ncbi.nlm.nih.gov/pubmed/28698636>.

73 Goncalves-Rodrigues, P., Almeida-Coelho, J., Goncalves, A., Amorim, F., Leite-Moreira, A. F., Stienen, G. J. M. & Falcao-Pires, I. In Vitro Assessment of Cardiac Function Using Skinned Cardiomyocytes. J Vis Exp, (2020). <https://www.ncbi.nlm.nih.gov/pubmed/32628167>.

74 Bell, G. I. Models for the specific adhesion of cells to cells. Science 200, 618–627, (1978). <https://www.ncbi.nlm.nih.gov/pubmed/347575>.

75 Kosuri, P., Alegre-Cebollada, J., Feng, J., Kaplan, A., Ingles-Prieto, A., Badilla, C. L., . . . Fernandez, J. M. Protein folding drives disulfide formation. Cell 151, 794–806, (2012). <https://www.ncbi.nlm.nih.gov/pubmed/23141538>.

76 Piotto, M., Saudek, V. & Sklenar, V. Gradient-tailored excitation for single-quantum NMR spectroscopy of aqueous solutions. J Biomol NMR 2, 661–665, (1992). <https://www.ncbi.nlm.nih.gov/pubmed/1490109>.

77 Wishart, D. S., Bigam, C. G., Yao, J., Abildgaard, F., Dyson, H. J., Oldfield, E., . . . Sykes, B. D. 1H, 13C and 15N chemical shift referencing in biomolecular NMR. J Biomol NMR 6, 135–140, (1995). <https://www.ncbi.nlm.nih.gov/pubmed/8589602>.

78 Delaglio, F., Grzesiek, S., Vuister, G. W., Zhu, G., Pfeifer, J. & Bax, A. NMRPipe: a multidimensional spectral processing system based on UNIX pipes. J Biomol NMR 6, 277–293, (1995). <https://www.ncbi.nlm.nih.gov/pubmed/8520220>.

79 Bartels, C., Xia, T. H., Billeter, M., Guntert, P. & Wuthrich, K. The program XEASY for computer-supported NMR spectral analysis of biological macromolecules. J Biomol NMR 6, 1–10, (1995). <https://www.ncbi.nlm.nih.gov/pubmed/22911575>.

80 Goddard, T. & Kneller, D. Sparky 3. University of California, San Francisco, USA. There is no corresponding record for this reference, (2008).

81 Improta, S., Politou, A. S. & Pastore, A. Immunoglobulin-like modules from titin I-band: extensible components of muscle elasticity. Structure 4, 323–337, (1996). <https://www.ncbi.nlm.nih.gov/pubmed/8805538>.

82 Tamiola, K. & Mulder, F. A. Using NMR chemical shifts to calculate the propensity for structural order and disorder in proteins. Biochem Soc Trans 40, 1014–1020, (2012). <https://www.ncbi.nlm.nih.gov/pubmed/22988857>.

83 Shen, Y., Delaglio, F., Cornilescu, G. & Bax, A. TALOS+: a hybrid method for predicting protein backbone torsion angles from NMR chemical shifts. J Biomol NMR 44, 213–223, (2009). <https://www.ncbi.nlm.nih.gov/pubmed/19548092>.

84 Hafsa, N. E., Arndt, D. & Wishart, D. S. CSI 3.0: a web server for identifying secondary and super-secondary structure in proteins using NMR chemical shifts. Nucleic Acids Res 43, W370–377, (2015). <https://www.ncbi.nlm.nih.gov/pubmed/25979265>.

85 Farrow, N. A., Muhandiram, R., Singer, A. U., Pascal, S. M., Kay, C. M., Gish, G., . . . Kay, L. E. Backbone dynamics of a free and phosphopeptide-complexed Src homology 2 domain studied by 15N NMR relaxation. Biochemistry 33, 5984–6003, (1994). <https://www.ncbi.nlm.nih.gov/pubmed/7514039>.

86 Dosset, P., Hus, J. C., Blackledge, M. & Marion, D. Efficient analysis of macromolecular rotational diffusion from heteronuclear relaxation data. J Biomol NMR 16, 23–28, (2000). <https://www.ncbi.nlm.nih.gov/pubmed/10718609>.

87 Lee, L. K., Rance, M., Chazin, W. J. & Palmer, A. G., 3rd. Rotational diffusion anisotropy of proteins from simultaneous analysis of 15N and 13C alpha nuclear spin relaxation. J Biomol NMR 9, 287–298, (1997). <https://www.ncbi.nlm.nih.gov/pubmed/9204557>.

88 Schrodinger, LLC. The PyMOL Molecular Graphics System, Version 1.*8* (2015).

89 Kroenke, C. D., Rance, M. & Palmer, A. G. Variability of the 15N Chemical Shift Anisotropy in Escherichia coli Ribonuclease H in Solution. Journal of the American Chemical Society 121, 10119–10125, (1999). <10.1021/ja9909273>.

90 Jarymowycz, V. A. & Stone, M. J. Fast time scale dynamics of protein backbones: NMR relaxation methods, applications, and functional consequences. Chem Rev 106, 1624–1671, (2006). <https://www.ncbi.nlm.nih.gov/pubmed/16683748>.

91 Akke, M., Brueschweiler, R. & Palmer, A. G., III. NMR order parameters and free energy: an analytical approach and its application to cooperative calcium(2+) binding by calbindin D9k. Journal of the American Chemical Society 115, 9832–9833, (1993). <10.1021/ja00074a073>.

92 Li, D. W. & Bruschweiler, R. A dictionary for protein side-chain entropies from NMR order parameters. J Am Chem Soc 131, 7226–7227, (2009). <https://www.ncbi.nlm.nih.gov/pubmed/19422234>.

93 Hopkins, J. B., Gillilan, R. E. & Skou, S. BioXTAS RAW: improvements to a free open-source program for small-angle X-ray scattering data reduction and analysis. J Appl Crystallogr 50, 1545–1553, (2017). <https://www.ncbi.nlm.nih.gov/pubmed/29021737>.

94 Konarev, P. V., Petoukhov, M. V., Volkov, V. V. & Svergun, D. I. ATSAS 2.1, a program package for small-angle scattering data analysis. Journal of Applied Crystallography 39, 277–286, (2006). <10.1107/S0021889806004699>.

95 Smith, S. B., Finzi, L. & Bustamante, C. Direct mechanical measurements of the elasticity of single DNA molecules by using magnetic beads. Science 258, 1122–1126, (1992). <https://www.ncbi.nlm.nih.gov/pubmed/1439819>.

