## Supplementary Information for "Pathological stiffening by crosslinking glycation of titin"

This file includes:

- Supplementary Figures S1-S15
- Supplementary Tables S1-S10
- Supplementary Text S1

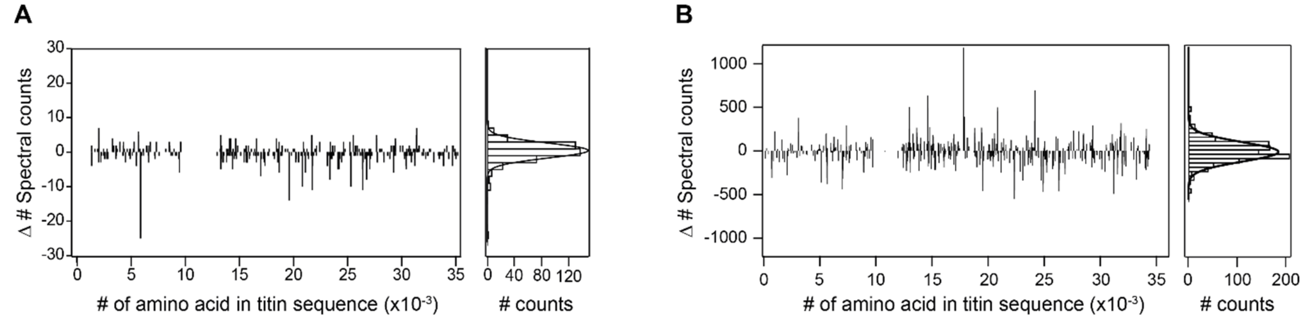


**Figure S1. Control distributions of spectral counts. (A)** To estimate the expected basal SD of the difference in the number of spectral counts (Δ#Spectral counts), the PSMs corresponding to peptidoforms modified at specific titin residues (on the x-axis) identified in 5 chromatographic runs from *ob/ob* and 5 from control mice were summed and subtracted from the sum of the equivalent peptidoforms identified in another 10 runs analyzed as technical replicates of the same samples. The global distribution of these differential events (right panel) was used to estimate the SD. **(B)** The same procedure was followed in the case of human native titin samples, by comparing two groups formed by one control and one diabetic myocardium sample.

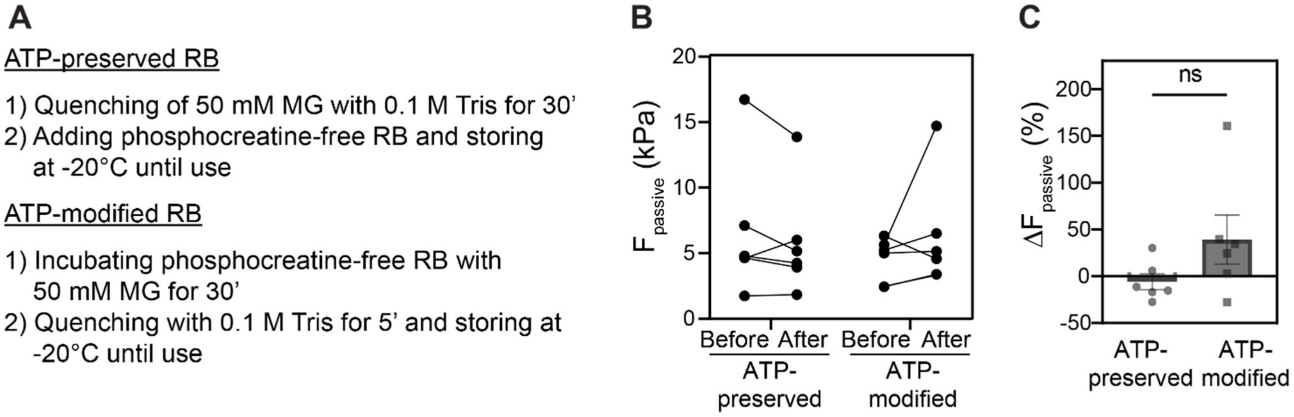


**Figure S2. Examination of the potential effect of MG-induced ATP modification on cardiomyocyte passive force. (A)** Scheme of the preparation of ATP-preserved and ATP-modified phosphocreatine-free relaxing buffer (RB). **(B)** Passive force values measured at 1.3 $L_0$ before and after the incubation with ATP-preserved and ATP-modified phosphocreatine-free RB (n = 6 cells per condition). **(C)** Changes in passive force measured at 1.3 $L_0$ after incubation with ATP-preserved and ATP-modified phosphocreatine-free RB. Error bars represent SEM. In B and C, each data point represents an individual cell. See Materials and Methods for details.

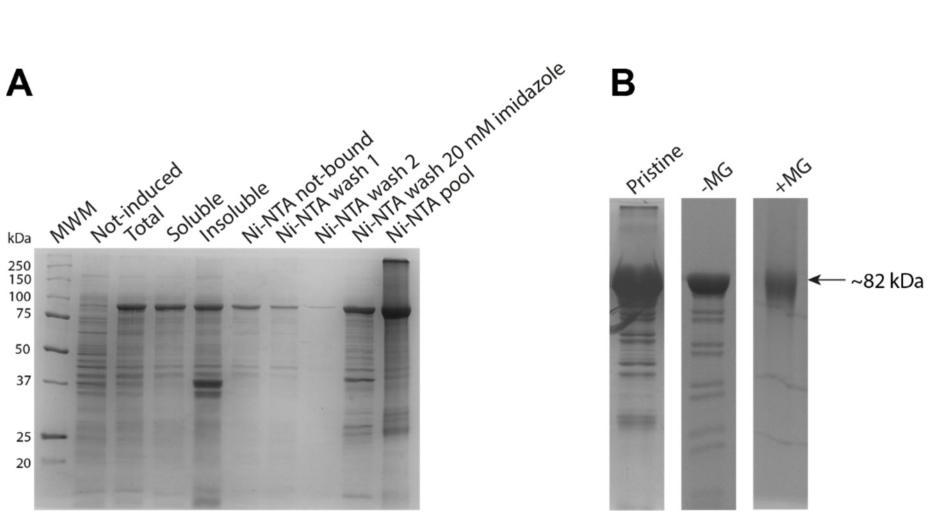


**Figure S3. (I91)$_8$ expression, purification and modification with MG. (A)** 12% SDS-PAGE gel to analyze protein expression and purification of (I91)$_8$. MWM: molecular weight markers (Precision Plus Protein™ Unstained Protein Standards, Bio-Rad). **(B)** 12% SDS-PAGE analysis of size-exclusion chromatography fractions at ~15 mL elution volume. Arrow marks the position of monomeric (I91)$_8$. Gels are stained with Coomassie blue.

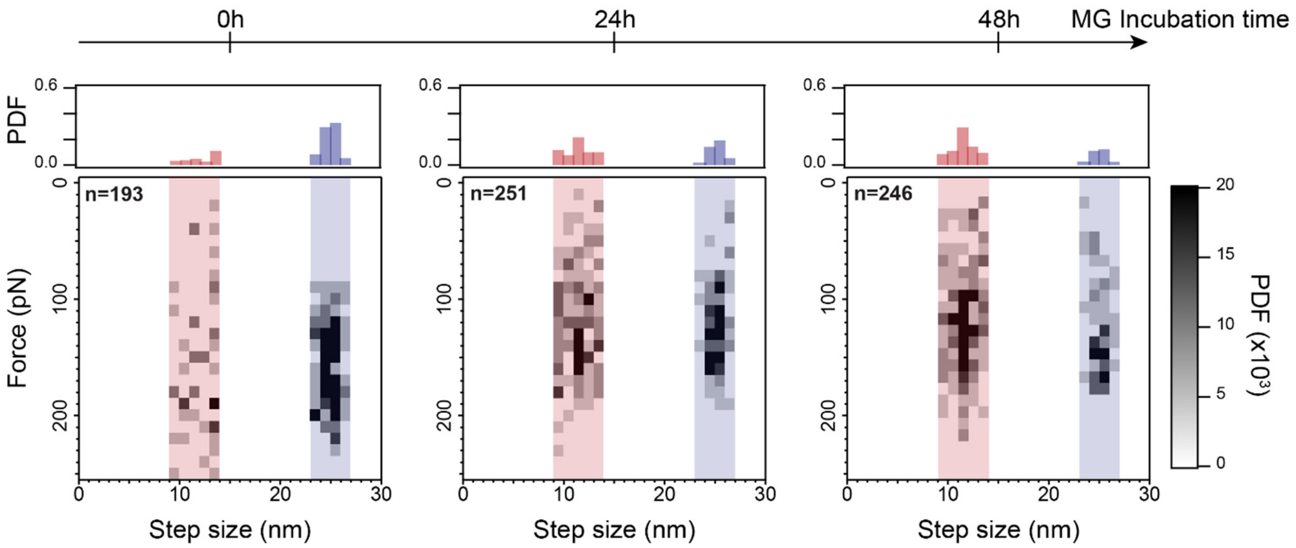


**Figure S4 Single-molecule analysis of $(I91)_8$ samples incubated with 50 mM MG for different times.** Data are presented as in **Figure 2K-M**.

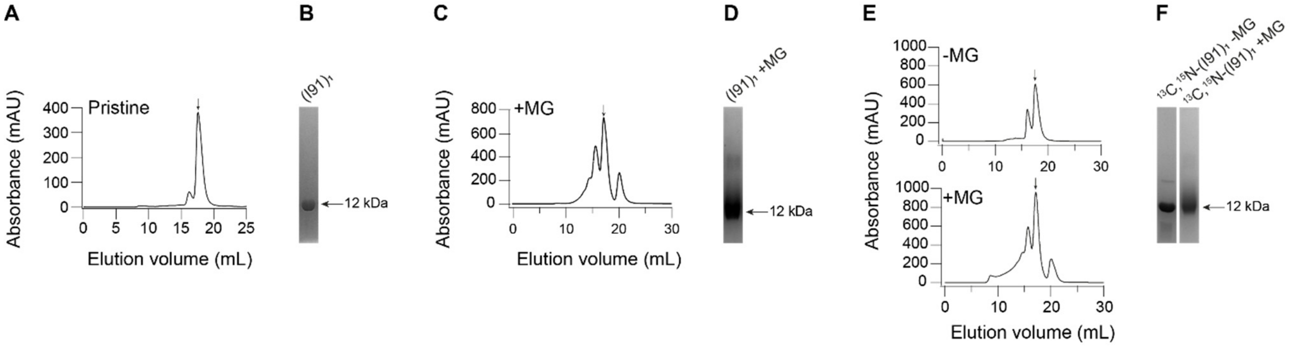


**Figure S5. Expression and purification of (I91)$_1$. (A, C, E)** Size-exclusion chromatograms of pristine (I91)$_1$ (A), glycated (I91)$_1$ (C) and $^{13}$C,$^{15}$N-(I91)$_1$ upon the incubation in the absence or presence of MG (E). Arrows mark the elution volume of monomeric I91. **(B, D, F)** 17% SDS-PAGE analysis of fractions with the highest absorbances. Gels are stained with Coomassie blue.

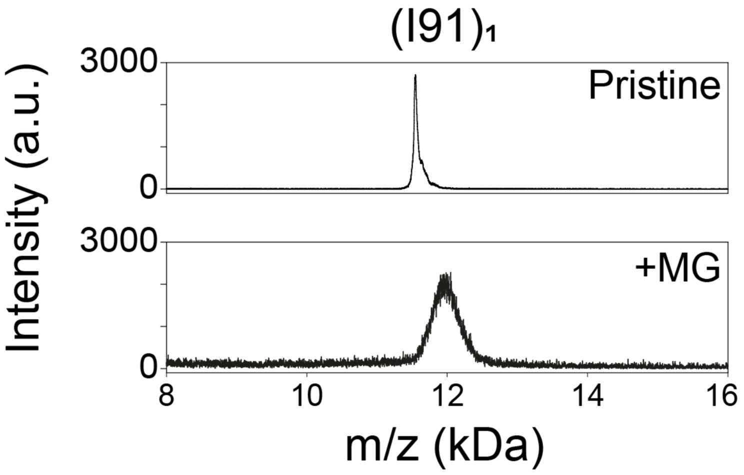


**Figure S6. MALDI-TOF/TOF spectra of pristine $(I91)_1$ and of the protein treated for 24 hours with 50 mM MG at 37°C.**

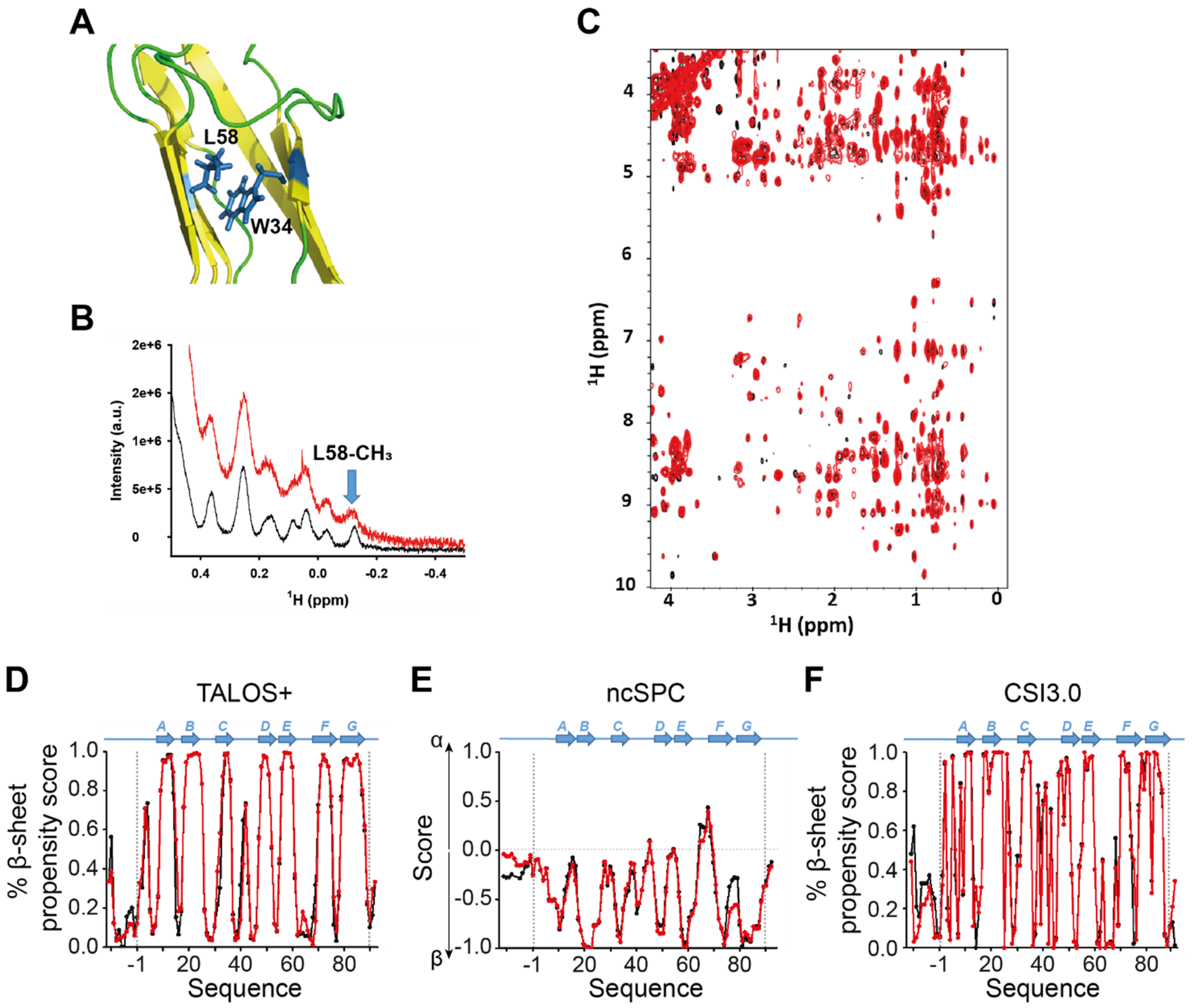


**Figure S7. Effect of MG treatment on the tertiary and secondary structure of $(I91)_1$. (A)** Ribbon representation of the contact region between L58 and W34 in the solution structure of I91 (PDB: 1TIT). Disordered regions are colored in green, whereas the β-sheet regions are colored in yellow. The side chains of L58 and W34 are shown as blue sticks. The image was built using Pymol[1]. **(B)** Overlapping of the high-field regions in the $^1H$ NMR spectra of non-glycated (black) and glycated (red) $(I91)_1$. The signal corresponding to the $–^{\delta}CH_3$ group of L58 is indicated with a blue arrow. **(C)** Overlapping of the projections corresponding to the $^1H$-$^1H$ planes of the $^{13}C$-NOESY-HSQC spectra of non-glycated (black) and glycated (red) $(I91)_1$. **(D)** β-sheet propensity scores obtained from TALOS+ [2] using the HN, N, $H_\alpha$, $C_\alpha$, $C_\beta$, and CO chemical shifts of non-glycated (black) and glycated (red) $(I91)_1$. **(E)** ncSPC scores obtained for non-glycated (black) and glycated (red) $(I91)_1$ calculated from the $H_N$, $H_\alpha$, $C_\alpha$, $C_\beta$ and CO chemical shifts. "+1" indicates the maximum propensity to form a full α-helix, "-1" indicates a fully formed β-sheet, and "0" indicates disordered region (indicated as a dashed gray horizontal line). **(F)** β-sheet propensity scores using the CSI3.0 software (http://csi3.wishartlab.com/cgi-bin/index.php) from the HN, N, $H_\alpha$, $C_\alpha$, $C_\beta$, and CO chemical shifts of non-glycated (black) and glycated (red) $(I91)_1$. In D-F, vertical dotted lines mark the beginning and end of the I91 domain (**Text S1**).

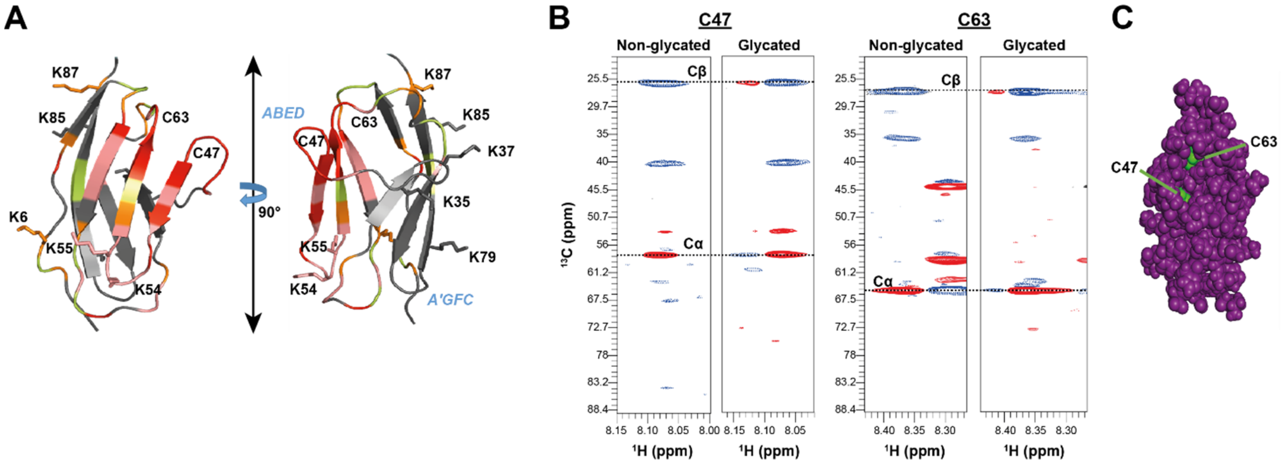


**Figure S8. Additional effects of glycation of $(I91)_1$ by NMR (A)** Carton representation of the 3D structure of I91 (PDB 1TIT), color-coded according to the $(I_{glyc}/I_{nat})$-1 values in **Figure 3H**. Red represents residues with values greater than 1.4, while salmon corresponds to values between 1.4 and 1.1. Orange is used for residues with values between 1.1 and 0.8, and lime green indicates those between 0.8 and 0.6. Residues with values below 0.6 are shown in gray, whereas cyan highlights those with values below 0. The image shows two 90°-rotated views of the same structure. The two beta-sheets of the I91 fold are indicated. **(B)** HNCACB strip plots at the $^{15}N$ frequencies for the amide resonances of C47 and C63 residues in the non-glycated and glycated $(I91)_1$. The $C_\alpha$ resonances are phased to yield positive peaks (red), whereas the HN-$C_\beta$ cross-peaks are negative (blue). The peaks corresponding to $C_\alpha$ and $C_\beta$ resonances are labeled. Glycation does not change the chemical shifts of $C_\alpha/C_\beta$ in C47 nor in C63. **(C)** Spacefill representation of the 3D structure of I91, where all the residues are colored in purple except C47 and C63, which are colored in green. Figure produced with Pymol[1]

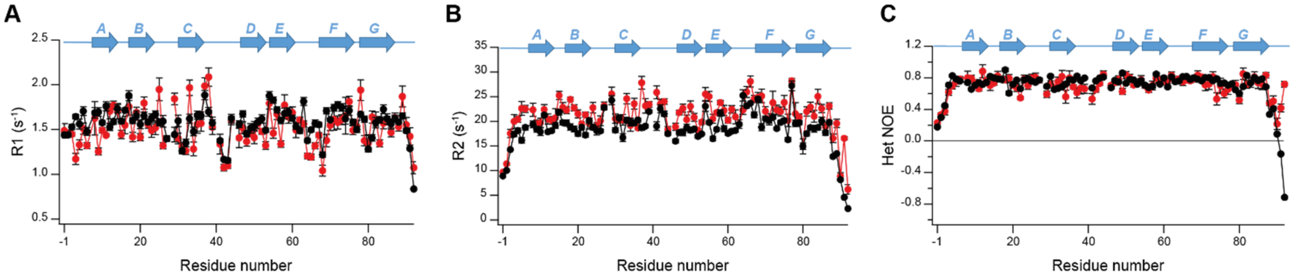


**Figure S9. Backbone NMR relaxation data**. Values of the backbone amide R1 (**A**), R2 (**B**) and $^{15}$N HET-NOE **(C)** obtained for non-glycated $(I91)_1$ (black) and glycated $(I91)_1$ (red). Location of β-strands is shown at the top of each panel.

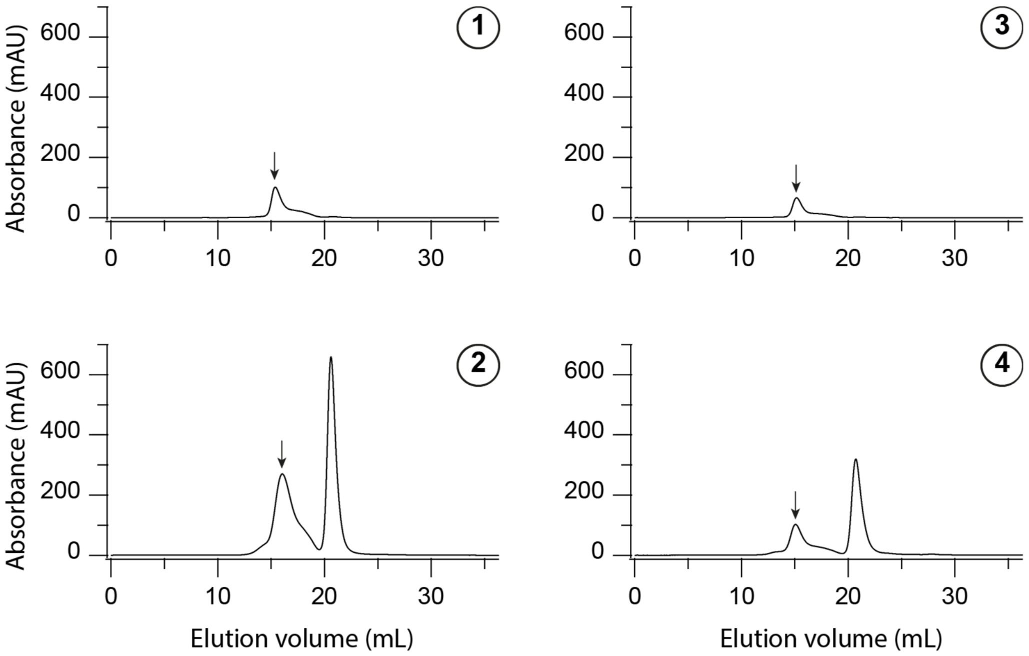


**Figure S10. Size-exclusion chromatograms corresponding to the $(I91)_8$ preparations according to the experimental design in Figure 4B.** Arrows indicate the elution volume of $(I91)_8$.

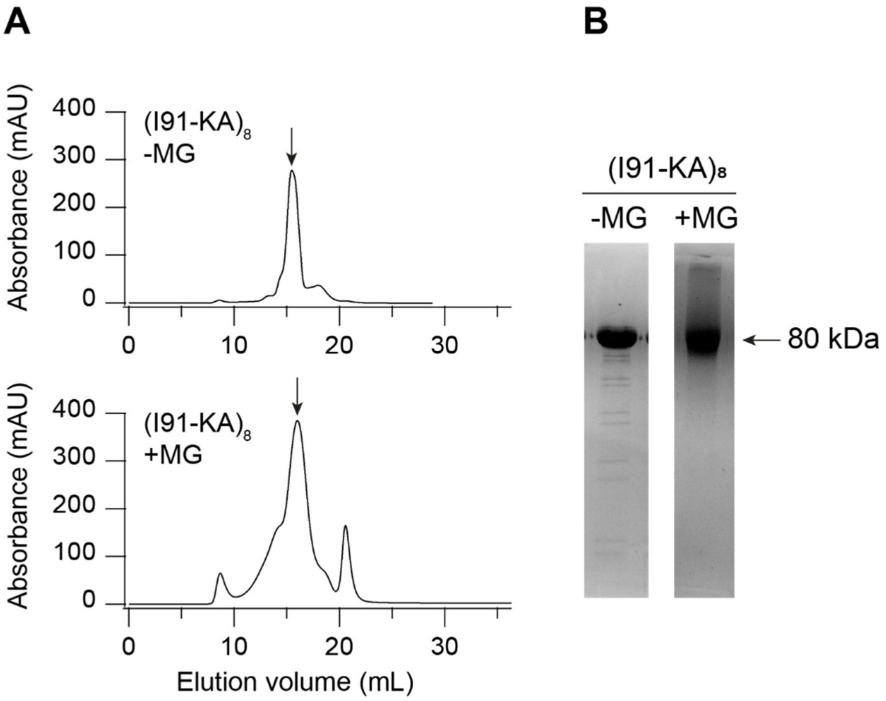


**Figure S11. Production of (I91-KA)$_8$. (A)** Size-exclusion chromatograms of (I91-KA)$_8$ after incubation in the absence or in the presence of MG. Arrows mark the elution volume of (I91-KA)$_8$. **(B)** 12% SDS-PAGE analysis of the fractions with the highest absorbance. Gels are stained with Coomassie blue.

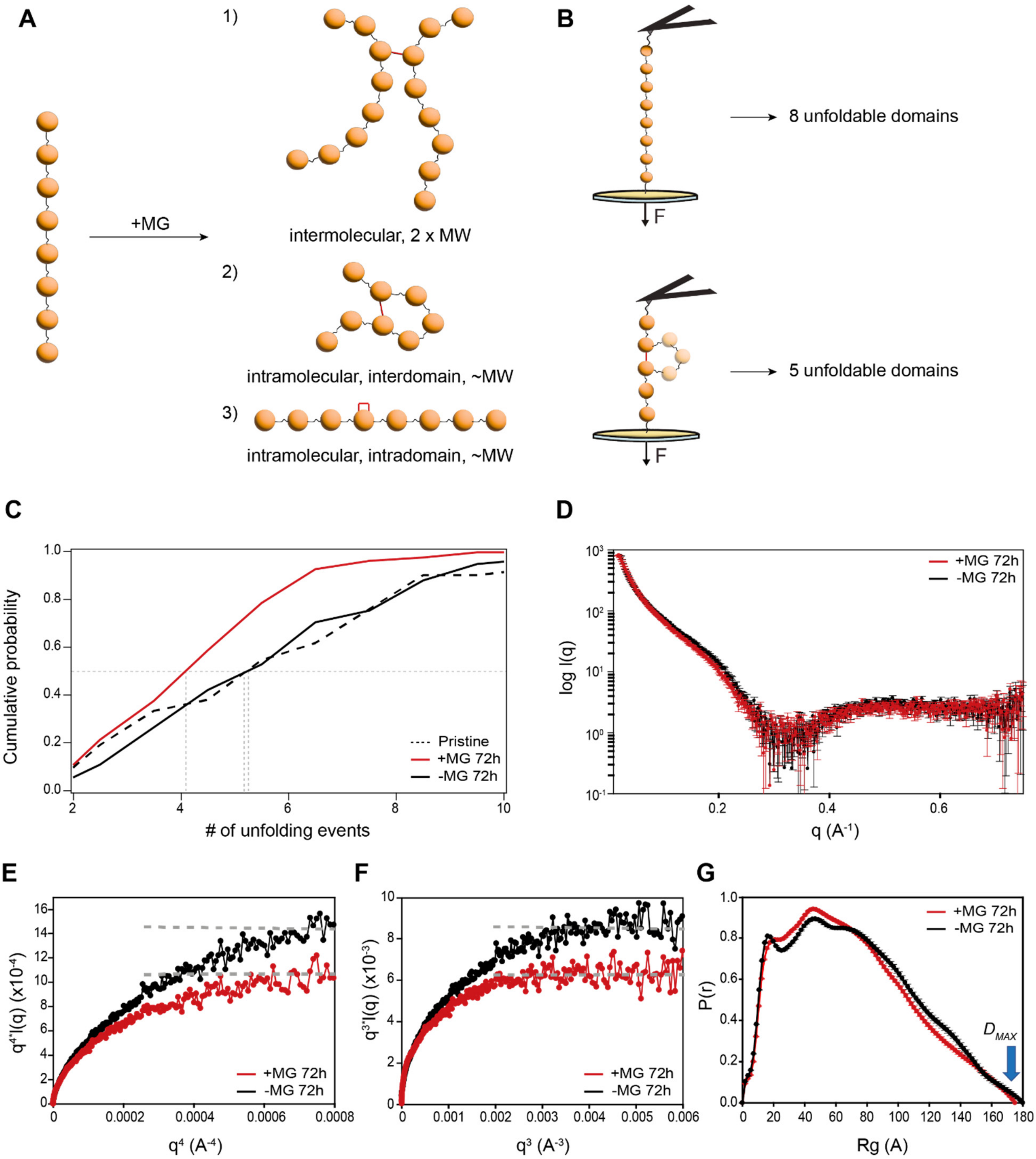


**Figure S12. Investigation of interdomain crosslinks. (A)** Scheme of different crosslinks that can be formed upon incubation of a polyprotein with MG. Crosslinks may be formed between different molecules (intermolecular, resulting in a molecule with double molecular weight, MW), but also within the same molecule (inter- or intradomain). **(B)** When a polyprotein does not contain crosslinks, all its domains can unfold in AFS experiments (*up*). In contrast, the presence of interdomain crosslinks prevents the protected domain from unfolding (*down*). **(C)** Cumulative probability of the number of unfolding events per trace for $(I91)_8$ preparations. **(D)** SAXS signal using a sample-detector distance of 370 mm obtained at 25ºC for $(I91)_8$ incubated in the presence (red) or absence (black) of MG for 72 h at 37ºC. **(E)** Porod-Debye plots obtained for $(I91)_8$ preparations. **(F)** Plot of the $q^3 \cdot I(q)$ values versus $q^3$ for the $(I91)_8$ preparations. **(G)** Pair distance distribution functions (P(r)) obtained for $(I91)_8$ preparations, based on the analysis of the experimental SAXS data using the program GNOM. The $D_{MAX}$ value obtained for both protein preparations is indicated by an arrow.

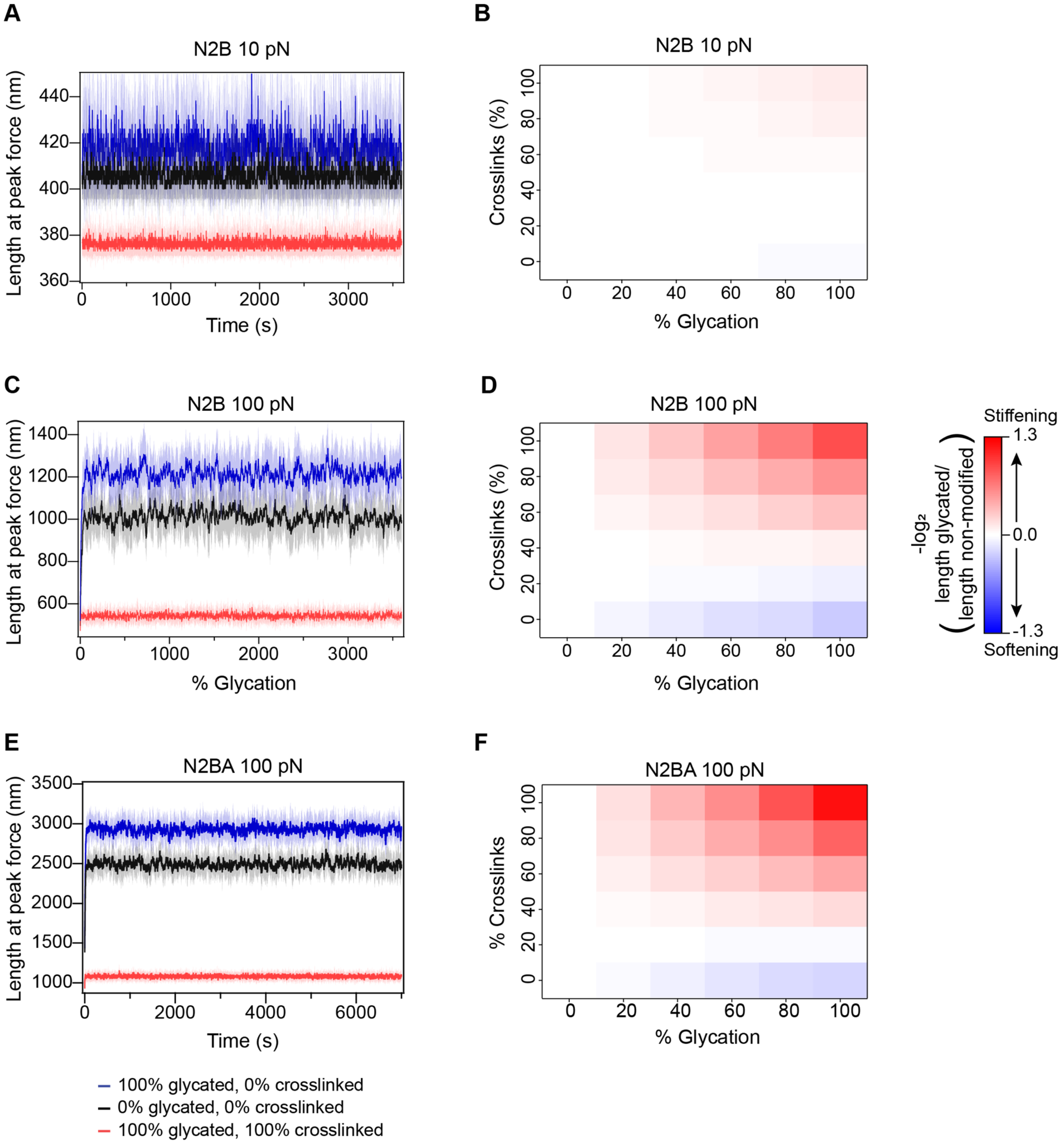


**Figure S13. Effects of glycation on titin mechanics studied by Monte Carlo simulations, considering PEVK domains are glycated.** **(A)** Length of N2B titin measured at 10 pN peak force during the simulations for non-modified (black), 100% glycated/0% crosslinked (blue) and 100% glycated/100% crosslinked (red) protein. Solid lines are the average of 10 independent simulations. Shaded areas represent SD. **(B)** Heatmap representing N2B titin length at 10 pN peak force for Monte Carlo simulations at different glycation conditions, relative to the non-modified protein. Results are the average of 10 independent simulations per condition. **(C)** Length of N2B titin measured at 100 pN peak force during the simulations for non-modified (black), 100% glycated/0% crosslinked (blue) and 100% glycated/100% crosslinked (red) protein. Solid lines are the average of 10 independent simulations. Shaded areas represent SD. **(D)** Heatmap representing N2B titin length at 100 pN peak force for Monte Carlo simulations at different glycation conditions, relative to the non-modified protein. Results are the average of 10 independent simulations per condition. **(E)** Length of N2BA titin measured at 100 pN peak force during the simulations for non-modified (black), 100% glycated/0% crosslinked (blue) and 100% glycated/100% crosslinked (red) protein. Solid lines are the average of 10 independent simulations. Shaded areas represent SD. **(F)** Heatmap representing N2BA titin length at 100 pN peak force for Monte Carlo simulations at different glycation conditions, relative to the non-modified protein. Results are the average of 10 independent simulations per condition.

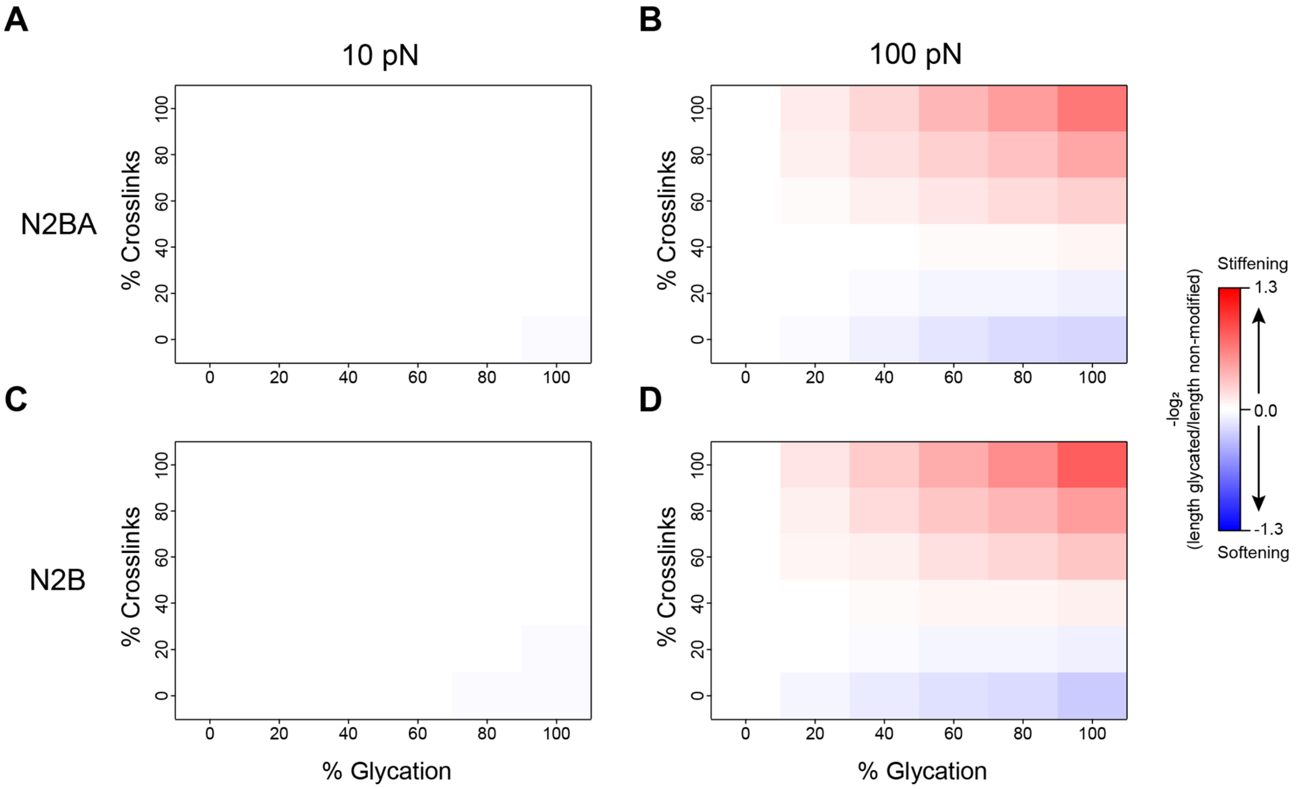


**Figure S14. Effects of glycation on titin mechanics studied by Monte Carlo simulations, considering PEVK domains are not glycated.** Heatmaps representing changes in titin length at different degrees of glycation for N2BA (A, B) and N2B (C, D) isoforms at 10 (A, C) and 100 (B, D) pN peak forces, not considering the possibility of glycation of the PEVK region, and comparing with non-modified protein. Results are the average of 10 independent simulations per condition.

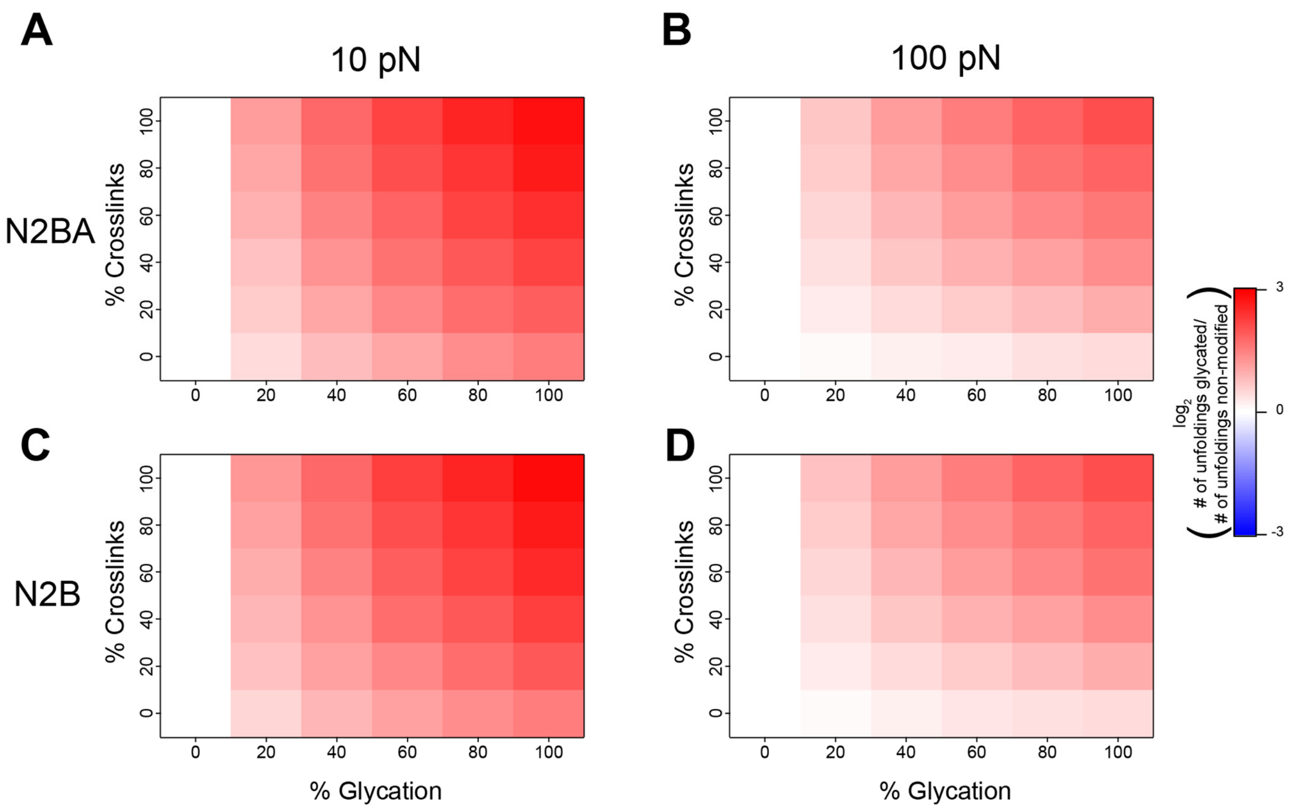


**Figure S15. Effects of glycation of titin folding/unfolding dynamics**. Heatmaps representing changes in number of unfolding events of titin isoforms in Monte Carlo simulations at different glycation conditions, compared with non-modified titin. Plots show results for N2BA (A, B) and N2B (C, D) titin isoforms at 10 (A, C) and 100 (B, D) pN peak forces. Results are the average of 10 independent simulations per condition.

**Table S1.** $^{1}H$- and $^{13}C$-NMR chemical shifts (ppm) of sMOLD collected at 600 MHz in 20 mM sodium acetate, pH 4.5 supplemented with 10% (v/v) $D_2O$. Atom numbers appear as displayed in **Figure 3C**.

| Atom number | $^{1}$H (ppm) | $^{13}$C (ppm) |
|---|---|---|
| **2** | 8.57 | 137.22 |
| **4** | 7.11 | 121.68 |
| **5** | - | 134.26 |
| **6** | 2.18 | 10.76 |
| **2’** | 4.01 | 48.7 |
| **3’** | 1.37, 1.27 | 23.68 |
| **4’** | 1.75 | 30.88 |
| **5’** | 1.76 | 32.29 |
| **6’** | 3.61 | 56.97 |
| **7’** | - | 177.1 |
| **2’’** | 4.03 | 51.35 |
| **3’’** | 1.32, 1.20 | 23.6 |
| **4’’** | 1.79 | 31.14 |
| **5’’** | 1.76 | 32.29 |
| **6’’** | 3.61 | 56.97 |
| **7’’** | - | 177.1 |

**Table S2.** Parameters obtained fitting the distributions of unfolding forces in **Figure 6C** to the Bell-Evans model. Errors correspond to 83% confidence intervals.

| Sample | $r_0$ ($s^{-1}$) | Δx (nm) |
|---|---|---|
| -MG 23-27 nm | 0.002 ± 0.001 | 0.16 ± 0.01 |
| +MG 9-14 nm | 0.016 ± 0.003 | 0.12 ± 0.01 |
| -CEL 23-27 nm | 0.007 ± 0.003 | 0.18 ± 0.02 |
| +CEL 23-27 nm | 0.021 ± 0.005 | 0.14 ± 0.01 |

**Table S3.** Mechanical parameters used for Monte Carlo simulations. Parameters were taken from[3,4], expect the reduction in contour length of unfolded Ig domains containing crosslinking AGEs, which is based on the results of this work.

| Element | Contour length (nm) | Kuhn length (nm) |
|---|---|---|
| PEVK (N2BA isoform) | 721 | 1.82 |
| PEVK (N2B isoform) | 68 | 1.82 |
| N2Bus | 230 | 1.32 |
| Folded Ig | 4 | 20 |
| Unfolded Ig (non-modified) | 35.2 | 1.32 |
| Unfolded Ig (non-crosslinking AGEs) | 35.2 | 1.32 |
| Unfolded Ig (crosslinking AGEs) | 18.8 | 1.32 |

**Table S4.** Transition rates for titin domain unfolding and refolding from the single-molecule AFS data in **Figure 6**. As AFM measurements cannot resolve the force-depending of folding, we chose the same Δx value for the three conditions according to published data[4,5]. For non-modified Ig domains, we used the parameters obtained for 191 incubated in the absence of MG for 72 h. Values for Ig domains containing crosslinking AGEs are those obtained experimentally in I91 samples after 72 h incubation with 50 mM MG. Values for domains containing non-crosslinking AGEs were obtained considering how the parameters in the CEL-containing sample change with respect to its control (the I91 sample incubated in the absence of pyruvic and $NaBH_3CN$), and adjusting accordingly the values to the non-modified domain parameters so that relative changes are maintained.

| | **Unfolding** | | **Folding** | |
|---|---|---|---|---|
| **Type of Ig domain** | **Δx (nm)** | **$r_0$ ($s^{-1}$)** | **Δx (nm)** | **$r_0$ ($s^{-1}$)** |
| Non-modified | 0.16 | $2.2 \times 10^{-3}$ | -2.2 | 4.6 |
| Containing crosslinking AGEs | 0.11 | $16.6 \times 10^{-3}$ | -2.2 | 29.7 |
| Containing non-crosslinking AGEs | 0.13 | $6.7 \times 10^{-3}$ | -2.2 | 4.6 |

**Table S5.** Folding fraction of non-glycated and glycated titin domains from fits in **Figure 6F,G**.

| Element | Non-modified | Crosslinking AGE | Non-crosslinking AGE |
|---|---|---|---|
| **Folding fraction** | 0.5 | 0.8 | 0.5 |

**Table S6.** Clinical characteristics of diabetic and non-diabetic patients.

| Sample | Age | Gender | DM | Years since diagnosis | Glycated hemoglobin (mg/dL) | Creatinin clearance (mL/min/m$^2$) | Hepatic disease | Cardiopulmonary disease | Oncohematological disease |
|---|---|---|---|---|---|---|---|---|---|
| A200016 | 51 | Male | No | - | - | 81.8 | No | No | No |
| A200017 | 74 | Male | Yes | 4 | 6.7 | 60 | No | No | No |
| A200018 | 71 | Male | No | - | - | >90 | No | No | Yes (Mantle cell lymphoma, stage IV A (gastric involvement and prostate adenocarcinoma, Gleason 8) |
| A200024 | 63 | Female | Yes | 7 | 7 | >90 | No | Yes (sleep apnea hypoventilation syndrome) | No |

**Table S7.** Composition of relaxing buffers (taken from [6]).

| Component | Concentration |
|---|---|
| **Imidazole-containing relaxing buffer (RB)** | |
| $Na_2ATP$ | 5.97 mM |
| $MgCl_2 \cdot 6H_2O$ | 6.04 mM |
| Tritiplex (EGTA) | 2 mM |
| KCl | 139.6 mM |
| Imidazole | 10 mM |
| **Propionic-containing relaxing buffer** | |
| $Na_2ATP$ | 5.89 mM |
| 1M $MgCl_2$ | 6.48 mM |
| Propionic acid | 40.76 mM |
| BES | 100 mM |
| Tritiplex (EGTA) | 6.97 mM |
| $Na_2PCr$ | 14.5 mM |
| **Phosphocreatine-free relaxing buffer** | |
| $Na_2ATP$ | 5.89 mM |
| 1M $MgCl_2$ | 6.48 mM |
| Propionic acid | 40.76 mM |
| BES | 100 mM |
| Tritiplex (EGTA) | 6.97 mM |
| NaCl | 43.5 mM |

**Table S8.** Composition of the isotopically labeled M9 minimal medium.

| Component | Concentration |
|---|---|
| $Na_2HPO_4$ | 33.7 mM |
| $KH_2PO_4$ | 22.0 mM |
| NaCl | 8.55 mM |
| $^{15}NH_4Cl$ | 18.7 mM |
| $^{13}C_6$-D-glucose | 0.2 % |
| $MgSO_4$ | 1 mM |
| $CaCl_2$ | 0.6 mM |
| Biotin | 1 µg/L |
| Thiamin | 1 µg/L |
| EDTA | 134 µM |
| $FeCl_3$-$6H_2O$ | 31 µM |
| $ZnCl_2$ | 6.2 µM |
| $CuCl_2$-$2H_2O$ | 0.76 µM |
| $CoCl_2$-$2H_2O$ | 0.42 µM |
| $H_3BO_3$ | 1.62 µM |
| $MnCl_2$-$4H_2O$ | 0.081 µM |

**Table S9.** Theoretical extinction coefficients values for monomeric and octameric I91 computed using ProtParam tool from ExPASy server[7].

| | **$(I91)_1$** | **$(I91)_8$** |
|---|---|---|
| **Extinction coefficients values ($M^{-1}$ $cm^{-1}$)** | 6990 | 55920 |

**Table S10. List of all possible I91 peptides crosslinked by MOLD considering proximity of lysine residues.** Peptide in bold type corresponds to the one identified in **Figure 5.** MH: monoprotonated monoisotopic mass. Numbers in the brackets (placed after crosslinked lysines) are the masses of MOLD plus the corresponding crosslinked peptides (indicated in the third column).

| MH | Sequence | Crosslinked sequence |
|---|---|---|
| 2934.503 | AASPDCEIIEDGKK[1038.55007]HIL | IEVEKPLY |
| 2771.503 | AASPDCEIIEDGKK[875.55007]HIL | IEVEKPL |
| 2658.503 | AASPDCEIIEDGKK[762.55007]HIL | EVEKPL |
| 2934.503 | IEVEK[1943.93117]PLY | AASPDCEIIEDGKKHIL |
| 2771.418 | IEVEK[1943.93117]PL | AASPDCEIIEDGKKHIL |
| 2658.334 | EVEK[1943.93117]PL | AASPDCEIIEDGKKHIL |
| 2529.292 | VEK[1943.93117]PL | AASPDCEIIEDGKKHIL |
| **1396.766** | **QAANTK[308.19637]SAANL** | **KL** |

**Text S1.** Sequence of the recombinant $(\mathrm{I91})_1$ protein. Extensions not present in titin are highlighted in gray. Lysines are highlighted in yellow.

MR$^{-11}$ GSHHHHHHGS$^{-1}$ L$^{1}$IEVEKPLYG$^{10}$ VEVFVGETAH$^{20}$ FEIELSEPDV$^{30}$ HGQWKLKGQP$^{40}$ LAASPDCEII$^{50}$ EDGKKHILIL$^{60}$ HNCQLGMTGE$^{70}$ VSFQAANTKS$^{80}$ AANLKVKELR$^{90}$ SC

### Supplementary references


1 Schrodinger, LLC. *The PyMOL Molecular Graphics System, Version 1.8* (2015).

2 Shen, Y., Delaglio, F., Cornilescu, G. & Bax, A. TALOS+: a hybrid method for predicting protein backbone torsion angles from NMR chemical shifts. *J Biomol NMR* **44**, 213-223, (2009). <https://www.ncbi.nlm.nih.gov/pubmed/19548092>.

3 Giganti, D., Yan, K., Badilla, C. L., Fernandez, J. M. & Alegre-Cebollada, J. Disulfide isomerization reactions in titin immunoglobulin domains enable a mode of protein elasticity. *Nat Commun* **9**, 185, (2018). <https://www.ncbi.nlm.nih.gov/pubmed/29330363>.

4 Li, H., Linke, W. A., Oberhauser, A. F., Carrion-Vazquez, M., Kerkvliet, J. G., Lu, H., . . . Fernandez, J. M. Reverse engineering of the giant muscle protein titin. *Nature* **418**, 998-1002, (2002). <https://www.ncbi.nlm.nih.gov/pubmed/12198551>.

5 Herrero-Galan, E., Martinez-Martin, I., Sanchez-Gonzalez, C., Vicente, N., Bonzon-Kulichenko, E., Calvo, E., . . . Alegre-Cebollada, J. Basal oxidation of conserved cysteines modulates cardiac titin stiffness and dynamics. *Redox Biol* **52**, 102306, (2022). <https://www.ncbi.nlm.nih.gov/pubmed/35367810>.

6 Goncalves-Rodrigues, P., Almeida-Coelho, J., Goncalves, A., Amorim, F., Leite-Moreira, A. F., Stienen, G. J. M. & Falcao-Pires, I. In Vitro Assessment of Cardiac Function Using Skinned Cardiomyocytes. *J Vis Exp*, (2020). <https://www.ncbi.nlm.nih.gov/pubmed/32628167>.

7 Wilkins, M. R., Gasteiger, E., Bairoch, A., Sanchez, J. C., Williams, K. L., Appel, R. D. & Hochstrasser, D. F. Protein identification and analysis tools in the ExPASy server. *Methods Mol Biol* **112**, 531-552, (1999). <https://www.ncbi.nlm.nih.gov/pubmed/10027275>.
